# C3a receptor signalling in ObRb-expressing neurons promotes leptin resistance and obesity-associated cardiac remodelling

**DOI:** 10.64898/2026.10.05.756676

**Authors:** Elias Rawish, Carolin Gragoll, Michael Schneider, Timo Rusack, Katharina Kurz, Violetta Pilorz, Sophia Marie Reiners, Sven Geisler, Janti HajAhmad, Nuria Lopez-Alcantara, Nada Alnaggar, Andreas Teufel, Mirko Otto, Jorge Duque Escobar, Thomas Stiermaier, Sven G. Meuth, Zouhair Aherrahrou, Henrik Oster, Walter Raasch, Tanja Zeller, Harald Langer, Jörg Köhl, Ingo Eitel

## Abstract

Obesity is associated with hypothalamic inflammation, leptin resistance and increased cardiometabolic risk, but the neuroimmune signals linking nutrient excess to these outcomes remain incompletely defined. Here we show that diet-induced obesity is accompanied by local complement activation in the mouse hypothalamus, leading to C3a generation that signals through C3a receptor (C3aR) on leptin receptor (ObRb)-expressing neurons and blunts leptin signaling. Deletion of C3aR in ObRb-expressing neurons preserves leptin responsiveness, reduces food intake and weight gain, and improves glucose homeostasis during dietary challenge. Systemic pharmacological C3aR targeting induces weight loss and improves leptin responsiveness in established obesity. Genetic or pharmacological C3aR targeting also attenuates obesity-associated diastolic dysfunction and myocardial fibrosis; pair-feeding experiments reveal an additional component of cardiac protection beyond reduced weight gain, associated with lower cardiac sympathetic innervation. In humans, circulating complement activation markers are elevated in obesity and decline after bariatric surgery. Together, our findings identify C3aR signaling in ObRb-expressing neurons as a neuroimmune pathway contributing to hypothalamic leptin resistance and obesity-associated cardiac remodeling, highlighting complement signaling as a potential therapeutic target in cardiometabolic disease.

## Introduction

Obesity is characterized by resistance to the anorexigenic actions of leptin acting primarily on neurons in the arcuate nucleus (ARC) of the hypothalamus to suppress food intake and control energy balance ^1–3^. Hypothalamic inflammation is a key component of this state ^2,4–6^. However, the molecular cues that translate local inflammatory stress into sustained failure of leptin receptor signalling and subsequent cardiometabolic end-organ dysfunction remain ill-defined. Because hypothalamic leptin pathways also regulate autonomic output^7^, such signals may couple disordered energy balance to obesity-associated cardiac remodelling. Complement mediators regulate glial activation, neuronal development, dendritic architecture and blood-brain barrier integrity in the central nervous system^8–11^. Although systemic complement activation and C3a receptor (C3aR) signalling have been linked to obesity in experimental and clinical settings^12–14^, a role for complement within hypothalamic feeding circuits, particularly in leptin-responsive neurons, has not been explored. Here, we investigated whether nutritional excess is associated with local complement activation and C3a generation in the hypothalamus and whether C3aR signalling in long form leptin receptor (ObRb)-expressing neurons contributes to impaired leptin responsiveness and obesity-associated cardiac remodelling. Using mouse genetics, pharmacological C3aR targeting and human complement profiling, we implicate neuronal C3aR signalling as a previously unknown and important contributor to leptin resistance and cardiometabolic remodelling.

## Results

### Nutritional stress is associated with hypothalamic complement activation

Previously, intracerebral C3a has been reported to stimulate feeding in rats^15^. These findings prompted us to assess whether complement is engaged locally within hypothalamic circuits that control energy balance. Wild-type (WT) mice fed a high-fat/high-fructose (HFHF) diet for 16 weeks developed the expected phenotype including increased adiposity, hyperleptinaemia, glucose intolerance, insulin resistance and blunted responses to exogenous leptin (Extended Data Fig. 1a-f). HFHF feeding increased immunoreactivity for activated C3 fragments within the ARC and C3a concentrations in whole hypothalamic lysates, consistent with enhanced local complement activation (Fig. 1a-c). Flow cytometric profiling of dissociated hypothalami, using established gating strategies^16^, showed expansion and activation of microglia together with increased astrocyte numbers (Fig. 1d-f), consistent with prior reports of hypothalamic inflammation during dietary excess^17^. Roughly half of microglia were C3-positive under both dietary conditions (Fig. 1g-h), but the proportion of C3-positive astrocytes increased on HFHF diet, consistent with reactive astrocytes representing a diet-responsive contributor to hypothalamic C3 (Fig. 1i-k). Together, these data indicate that obesity is associated with local complement activation in hypothalamic regions involved in appetite regulation.

**Fig. 1.**
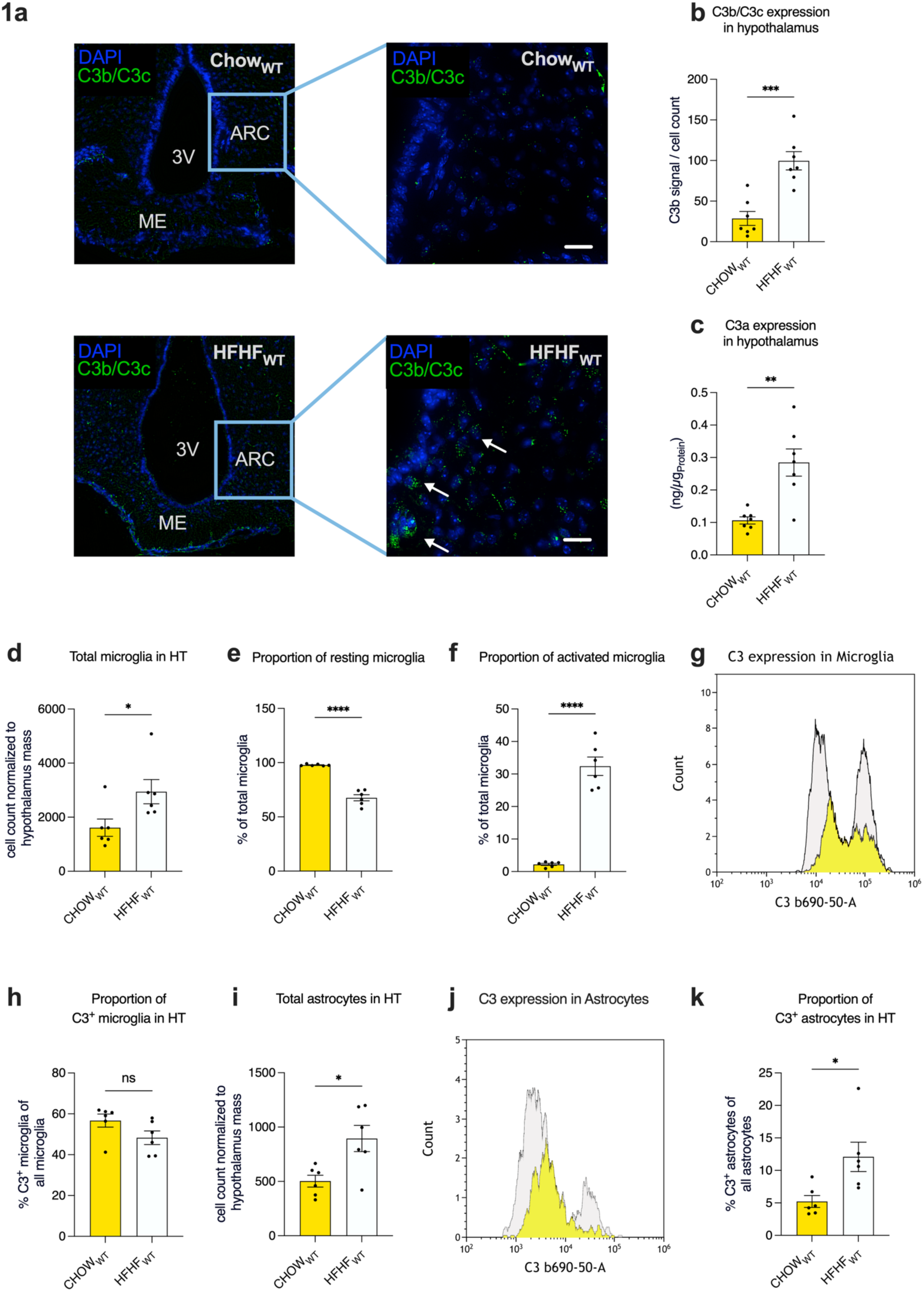
High-fat/high-fructose diet induces hypothalamic complement activation and glial reactivity associated with increased astrocytic C3 production. a, Immunohistochemical assessment of activated C3 fragment immunoreactivity in the arcuate nucleus (ARC) of CHOW-(top) or HFHF-fed (bottom) WT mice on day 112; white arrows point toward C3b/iC3b/C3c deposition (green). Scale bar: 25 µm b, Quantification of (i)C3b/C3c deposition in the ARC of HFHF- and CHOW-fed mice (n=7/group). c, C3a expression in the hypothalami of HFHF- and CHOW-fed mice on day 112 (n=7/group). d-f, Number (d) and frequency (e) of resting (CD11b⁺CD45^low^CD68^low^) or (f) activated microglia (CD11b⁺CD45^int^CD68^high^) in the hypothalami of HFHF- and CHOW-fed mice on day 112 (n=6/group). g, Representative flow cytometric histogram of C3 expression in hypothalamic microglia from HFHF (grey)- or CHOW-fed mice (yellow). h, Percentage of C3^+^ microglia (n=6/group). i, Number of astrocytes (GFAP^+^) in the hypothalami of HFHF- or CHOW-fed mice on day 112 (n=6/group). j, Representative flow cytometric histogram of C3 expression in hypothalamic astrocytes from HFHF (grey)- or CHOW-fed mice (yellow). k, Percentage of C3^+^ astrocytes (n=6/group). Data are shown as mean ± SEM. Statistical differences between groups were determined using unpaired Student’s t-test (b-f, h-i, k). ns: not significant, *p<0.05, **p<0.01, ***p<0.001, ****p<0.0001.

### C3aR signalling attenuates leptin responses in hypothalamic neurons

We next asked whether C3a can directly act on leptin-responsive hypothalamic neurons. First, we used the murine hypothalamic neuronal line mHypoE-N46, which expresses orexigenic agouti-related peptide (AgRP) and the long leptin receptor isoform ObRb. Importantly, mHypoE-N46 also expressed C3aR, which colocalized with ObRb (Fig. 2a-b). Stimulation with interleukin-6 (IL-6), a cytokine that intensifies hypothalamic inflammation^18^, increased C3aR expression, suggesting that inflammatory signals upregulate this receptor in hypothalamic neurons (Fig. 2c). In vivo, tdTomato C3aR reporter mice^19^ confirmed receptor expression in hypothalamic neurons in addition to microglia (Fig. 2d-e). To assess whether C3aR is expressed in leptin-responsive neurons, we administered leptin in vivo to tdTomato-C3aR mice, followed by analysis of phosphorylated STAT3 (pSTAT3) in the hypothalamus, since leptin receptor signalling depends on JAK2–STAT3 activation^20^. Leptin-induced pSTAT3 immunoreactivity co-localized with a subset of C3aR-positive neurons, consistent with C3aR expression in leptin-responsive hypothalamic neurons in vivo (Fig. 2f).

**Fig. 2.**
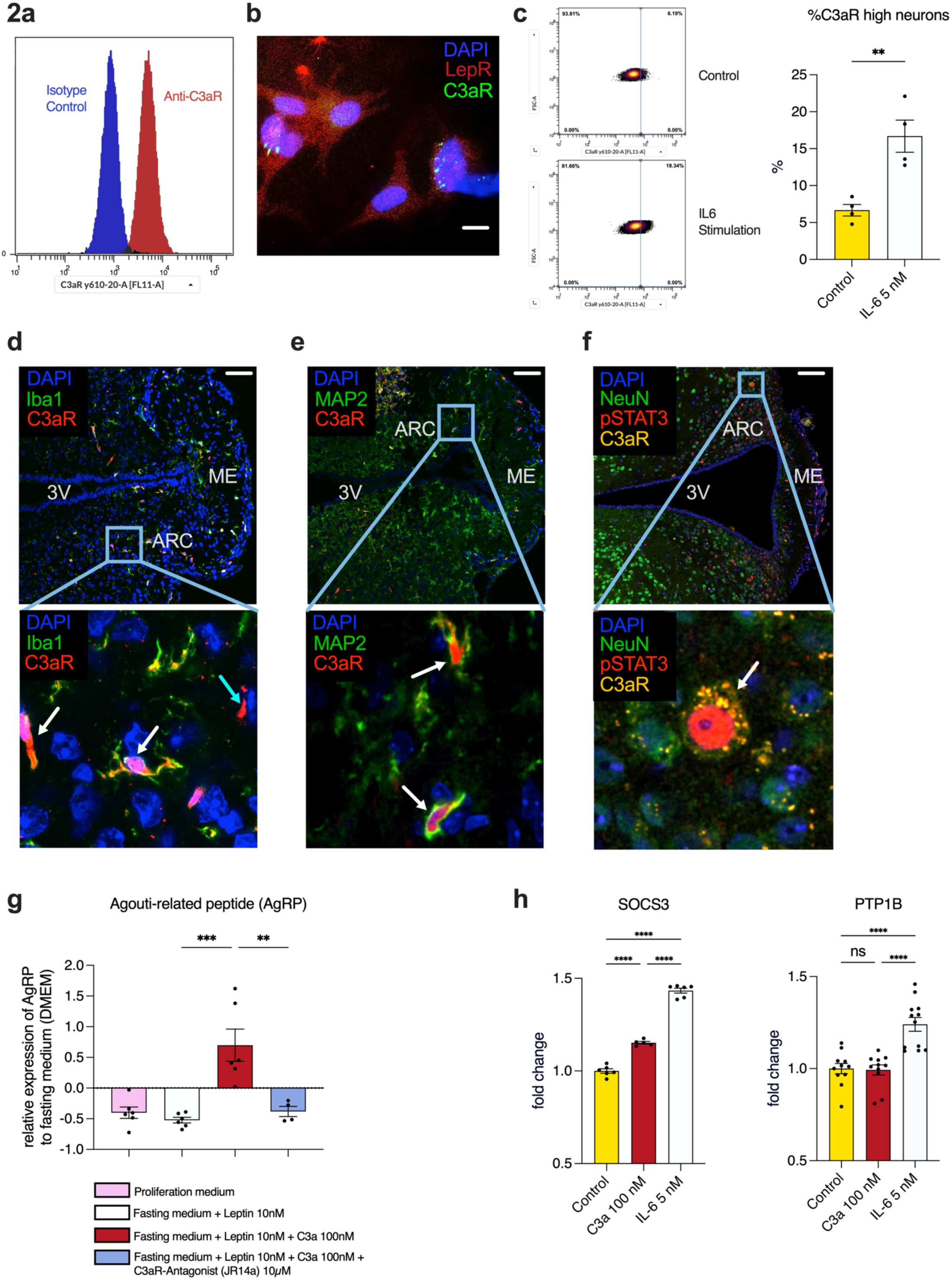
C3aR expression and regulation in hypothalamic neurons and its impact on leptin signaling. **a**, Representative histogram from flow cytometric analysis of mHypoE-N46 neurons stained with anti-C3aR antibody (green) versus isotype control (grey). **b**, Immunocytochemical staining of mHypoE-N46 neurons showing co-localization of leptin receptor (ObRb, red) and C3aR (green); nuclei are counterstained with DAPI (blue). Scale bar: 10 µm. **c**, Impact of IL-6 stimulation on C3aR expression in mHypoE-N46 neurons. Cells were treated with IL-6 for 12 hours. **Left:** Representative dot-plot showing C3aR expression in the absence of IL-6 (control) and after IL-6 stimulation. **Right:** Frequency of C3aR^high^ neurons in the absence of IL-6 and after IL-6 stimulation. **d**, Representative immunohistochemical image from C3aR-tdTomato-floxed reporter mice, showing co-localization of C3aR (red) with Iba1-positive microglia (green) in the arcuate nucleus (ARC) marked by white arrows; the cyan arrow points toward a C3aR-positive, Iba1-negative cell, suggesting expression in non-microglial cell types. ME = median eminence, 3V = third ventricle. Scale bar: 100 µm. **e**, Representative immunohistochemical image of hypothalamic sections showing co-expression of C3aR (red) with the neuronal marker MAP2 (green), confirming neuronal expression of C3aR in vivo. Scale bar: 100 µm. **f,** Representative immunofluorescence image of the hypothalamus from leptin-stimulated mice, showing pSTAT3-positive neurons (red) co-localizing with NeuN (green) and C3aR (yellow). pSTAT3 marks leptin-responsive neurons, demonstrating that C3aR is expressed in functionally leptin-responsive neuronal populations. Scale bar, 100 µm. **g**, Relative AgRP expression in mHypoE-N46 cells in response to leptin in the presence or absence of C3a and the C3aR modulator JR14a, normalized to fasting control (DMEM without glucose medium) (n=4–6/group). **h,** mRNA expression of SOCS3 (left) and PTP1B (right) in response to C3a or IL-6 (n=5–12/group) compared to PBS control. Cells were cultured in proliferation medium. Data shown are the mean ± SEM. Statistical differences between groups were assessed using Mann–Whitney U-test (**c**) or one-way ANOVA with Tukey’s multiple comparisons test (**g, h**). ns: not significant, **p<0.01, ***p<0.001, ****p<0.0001.

Functionally, C3a attenuated leptin-mediated suppression of AgRP in mHypoE-N46 cells. JR14a, a thiophene C3aR modulator originally described as an antagonist of C3a-induced C3aR responses^21,22^, prevented this C3a effect, supporting C3aR-dependent pharmacological engagement in this assay (Fig. 2g). Next, we tested whether C3aR signalling intersects with canonical leptin inhibitory programs. C3a induced SOCS3, a central negative regulator of leptin signalling^2,6^, whereas the insulin signaling pathway regulator protein tyrosine phosphatase 1B (PTP1B) remained unchanged (Fig. 2h), arguing for a selective intersection with the leptin– STAT3 feedback axis rather than broad activation of leptin resistance programs. Together, these findings support a model in which locally generated C3a can act directly on hypothalamic neurons to blunt leptin responsiveness.

### C3aR signalling in ObRb-expressing neurons promotes diet-induced obesity

To test the in vivo relevance of this pathway, we generated mice lacking C3aR in leptin receptor-expressing neurons (tdTomato-C3aR^flox/flox^ x ObRb-Cre). The long leptin receptor isoform ObRb is highly enriched in the hypothalamus^23,24^, but low-level Cre activity has been reported outside the brain in bone marrow mesenchymal stem cells (MSCs) in some settings^25^. Reporter analysis confirmed loss of neuronal C3a receptor signal in the hypothalamus, whereas microglial expression was preserved and no expression on hypothalamic astrocytes was observed (Fig. 3a–b, Extended Data Fig. 4a). In the peripheral and non-hypothalamic brain compartments examined, we did not detect evidence of substantial recombination affecting C3aR expression in piriform cortex, blood, bone marrow, heart or peripheral metabolic tissues^19^ (Extended Data Figs. 2, 3 and 4d-e, Supp. Table 1). Under chow feeding, body weight gain in control and conditional C3aR knockout mice were indistinguishable (Extended Data Fig. 4c). However, neuronal C3aR deficiency markedly limited body-weight gain and gonadal fat accumulation, with divergence emerging early during dietary challenge in response to HFHF feeding (Fig. 3c-d).

**Fig. 3.**
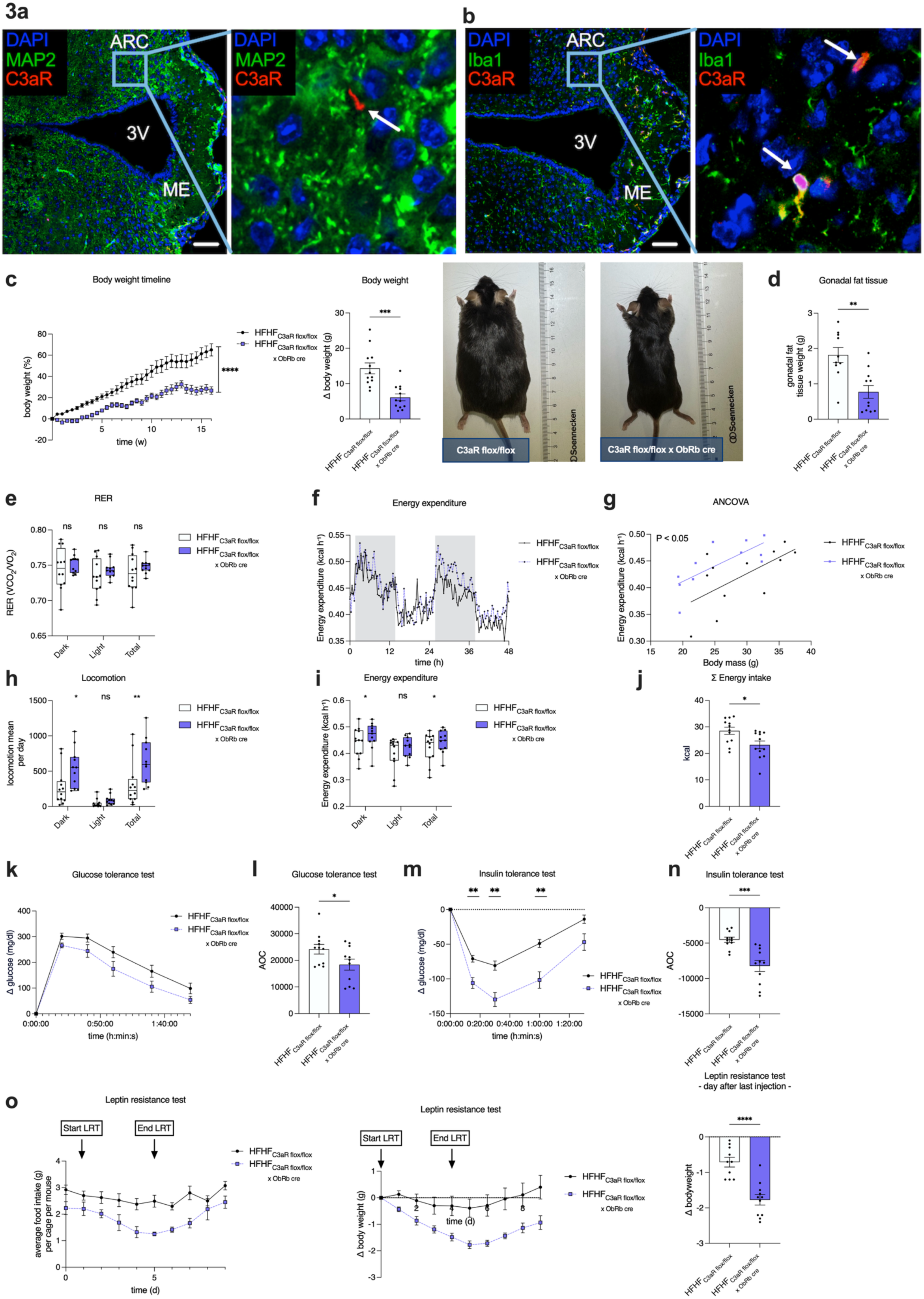
Deletion of C3aR in ObRb-expressing neurons limits diet-induced weight gain and improves metabolic responses. **a**, Representative immunohistochemical staining of hypothalamic sections from C3aR^flox/flox^ x ObRb-Cre mice. Co-staining of MAP2 (green), and tdTomato-labeled C3aR (red) shows no co-localization of MAP2 and tdTomato-C3aR, consistent with reduced tdTomato-C3aR signal in MAP2-positive hypothalamic neurons within the analysed region. DAPI is stained in blue. The white arrows point toward a C3aR^+^ cell that does not express MAP2. ARC = arcuate nucleus, ME = median eminence, 3V = third ventricle. Scale bar: 100 µm. **b**, Immunohistochemical staining of Iba1 (green) and tdTomato-labeled C3aR (red), in the hypothalamus of the conditional C3aR knockout mice indicates preserved microglial tdTomato-C3aR signal. White arrows point toward microglia-expressing tdTomato-C3aR. Scale bar: 100 µm. **c, Left:** Relative body weight gain of conditional C3aR-deficient and control mice fed a HFHF diet over 16 weeks, expressed as percentage change from baseline weight (n=12/group). **Middle:** Comparison of body weight gain in conditional C3aR-deficient vs. tdTomatoC3aR^flox/flox^ controls at week 16; shown is the relative increase to baseline weight (n=12/group). **Right:** Representative images of conditional C3aR-deficient and control mice after HFHF exposure at week 16. **d,** Comparison of gonadal white adipose tissue mass in conditional C3aR-deficient vs control mice at week 16 (n=10-11/group). **e,** Respiratory exchange ratio (RER) at week 10 measured by indirect calorimetry. Quantification during dark and light phases and across the total recording period (box plots show median and interquartile range; whiskers, minimum to maximum). **f,** Energy expenditure at week 10 measured by indirect calorimetry. **g,** ANCOVA for energy expenditure with body mass as a covariate. The groups are statistically different (p < 0.05), with no mass × group interaction. **h,** Locomotion under standard housing conditions. Quantification during dark and light phases and across the total recording period (box plots as in **e**). **i,** Energy expenditure quantified during dark and light phases and across the total recording period (box plots as in **e**). **j**, Energy intake (kcal) at week 10 in conditional C3aR-deficient and control mice measured by indirect calorimetry; n=11 conditional C3aR KO, n=12 control in **e-j**. **k,** Baseline-subtracted glucose levels during glucose tolerance test (GTT) at week 12 (n=11/group). **l,** Area over the curve (AOC) from GTT calculated from baseline-subtracted glucose levels (n=11/group). **m,** Baseline-subtracted glucose levels during insulin tolerance test (ITT) at week 13 (n=11/group). **n,** AOC from ITT calculated from baseline-subtracted glucose levels (n=11/group). **o, Left:** Daily food intake (g/d) during leptin resistance test (LRT) at week 14 (n=11/group). **Middle:** Body weight changes relative to values prior to LRT start (n=11/group). **Right:** Body weight change assessed the day after the final leptin injection in the LRT (n=10-11/group). Data shown are the mean ± SEM. Statistical differences between groups were determined using two-way ANOVA with Bonferroni’s multiple comparisons test (**c**, left; **e, h-i, m**), ANCOVA (**g**) or unpaired Student’s t-test (**c**, middle; **d, j, l, n, o**, right). ns: not significant, *p<0.05, **p<0.01, ***p<0.001., ****P<0.0001.

**Fig. 4.**
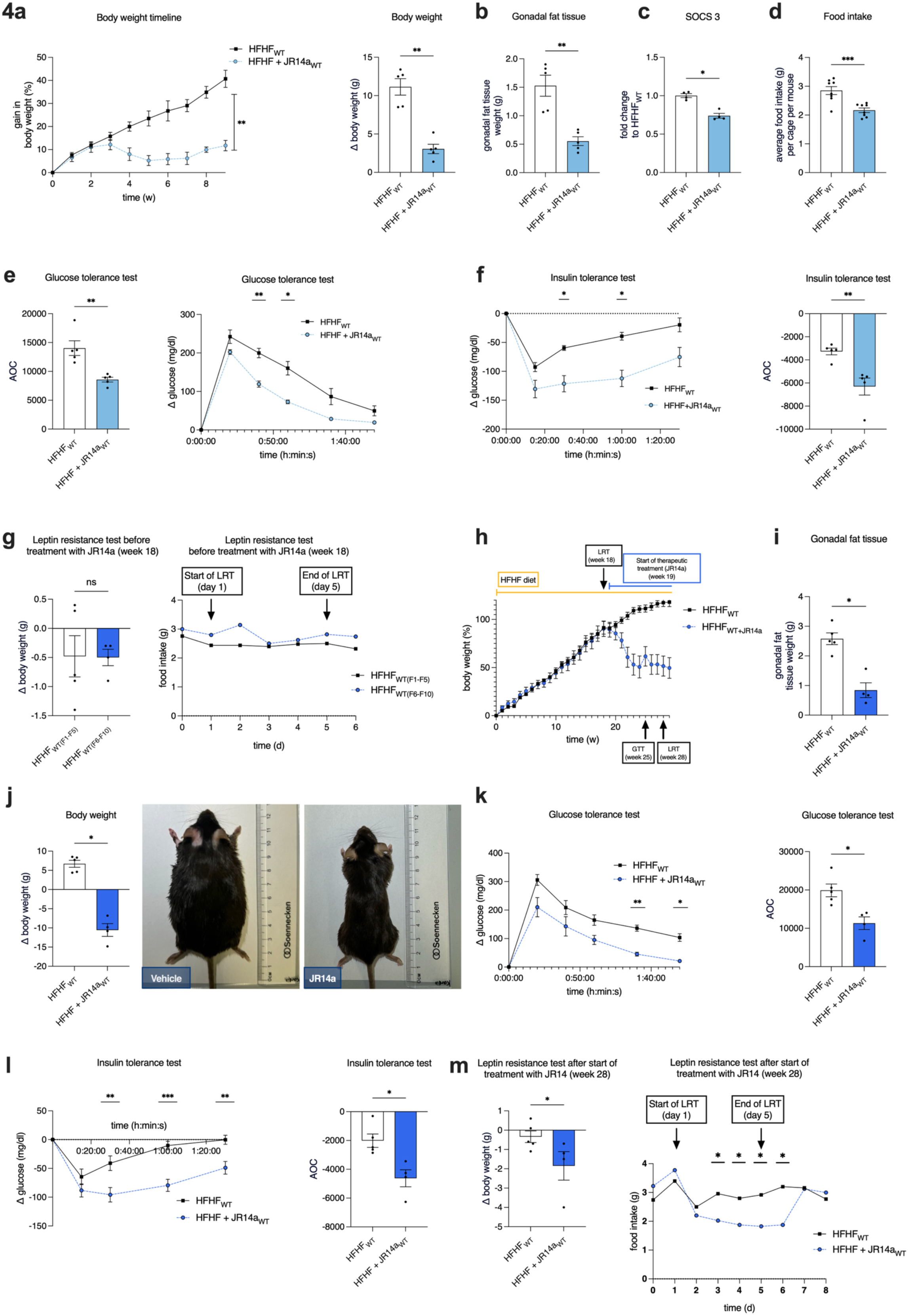
Preventive and therapeutic pharmacological C3aR targeting markedly reduce body weight gain in response to HFHF and improve insulin and leptin sensitivity. **a**, Top left: Impact of preventive C3aR targeting by JR14a on relative body weight gain in WT mice in response to HFHF diet over 9 weeks, expressed as percentage change from baseline weight (n=5/group). Top right: Body weight gain at week 9 relative to baseline weight (n=5/group). b, Mass of gonadal white adipose tissue at week 9 (n=5/group). c, SOCS3 mRNA expression in hypothalami at week 9 (n=4/group). d, Average daily food intake per mouse during week 7 (n=8/group). e, Left: Baseline-subtracted glucose levels during GTT (n=5/group). Right: AOC from GTT performed at week 8, calculated from baseline-subtracted glucose levels (n=5/group). f, Left: Baseline-subtracted glucose levels during ITT (n=5/group). Right: AOC from ITT performed at week 9, calculated from baseline-subtracted glucose levels (n=5/group). g, Left: Body weight change after the final leptin injection in LRT at week 18 before therapeutic intervention (n=4-5/group). Right: Quantification of food intake changes after leptin administration per cage per mouse. h, Representative experimental timeline and relative body weight change normalized to baseline in WT-mice treated with either vehicle or C3aR modulator JR14a (n=4 JR14a-treated group, n=5 control). i, Mass of gonadal white adipose tissue at week 30 (n=4 JR14a-treated group, n=5 control). j, Left: Body weight change (g) after the start of therapeutic treatment with JR14a or vehicle at week 30. (n=4 JR14a-treated group, n=5 control). Right: Representative images of WT-mice receiving vehicle or JR14a at week 30. **k, Left:** Baseline-subtracted glucose levels during GTT (n=4 JR14a-treated group, n=5 control group). **Right:** AOC from GTT performed at week 25 (n=4 JR14a-treated group, n=5 control). **l, Left:** Baseline-subtracted glucose levels during ITT (n=4 JR14a-treated group, n=5 control). **Right:** AOC from ITT performed at week 26 (n=4 JR14a-treated group, n=5 control). **m, Left:** Body weight change assessed the day after the final leptin injection at week 28 in the LRT after the start of therapeutic intervention (n=4 JR14a-treated group, n=5 control). **Right:** Quantification of food intake changes after leptin administration per cage per mouse. Data shown are the mean ± SEM. Statistical differences between groups were assessed using two-way ANOVA with Bonferroni’s multiple comparisons test (**a**, top left; **e**, left; **f**, left; **k**, left; **l**, left; m, left), Mann–Whitney U-test (**a**, top right; **b-c**; **e**, right; **f**, right; **g**, left; **i, j**, left; **k**, right; **l**, right; **m**, right) or unpaired Student’s t-test (**d**). ns: not significant, *p<0.05, **p<0.01, ***p<0.001.

Further, indirect calorimetry revealed lower cumulative food intake together with increased dark-phase energy expenditure and locomotor activity, without a change in respiratory exchange ratio (Fig. 3e-j). C3aR conditional knockout mice also showed improved glucose tolerance and insulin sensitivity and, importantly, retained robust anorectic and weight-reducing responses to leptin despite HFHF feeding (Fig. 3k-o). Thus, C3aR signalling in leptin receptor neurons is dispensable for basal energy homeostasis but becomes an important contributor to hyperphagia, leptin resistance and metabolic dysfunction during nutritional excess.

### Pharmacological C3aR targeting attenuates established obesity

To determine the translational potential of this finding, we pharmacologically targeted C3aR as an orthogonal validation strategy. First, we administered JR14a at the beginning of HFHF feeding. This treatment suppressed weight gain, reduced adiposity, lowered hypothalamic SOCS3 expression and improved glucose homeostasis (Fig. 4a-c). Food intake was also reduced, consistent with improved leptin responsiveness (Fig. 4d).

Next, we targeted C3aR after 18 weeks of HFHF feeding, when WT mice were overtly obese and leptin resistant (Fig. 4g-h). JR14a treatment induced progressive weight loss, reduced gonadal fat mass and improved leptin responsiveness, as reflected by greater leptin-associated suppression of body weight during challenge (Fig. 4h-j). Glucose tolerance, insulin sensitivity and leptin resistance also improved significantly (Fig. 4k-m). The concordance between neuron-specific genetics and systemic pharmacology suggests that C3aR signalling contributes not only to the development of obesity but also to the maintenance of the leptin-resistant state.

### C3aR signalling in ObRb-expressing neurons contributes to obesity-associated cardiac remodelling

Because obesity-associated leptin resistance is increasingly linked to cardiometabolic end-organ disease^26–28^, we examined cardiac structure and function. After 16 weeks of HFHF diet, floxed control mice exhibited echocardiographic features consistent with diastolic dysfunction, including increased left-ventricular wall thickness, higher E/E′ and prolonged isovolumic relaxation time, while ejection fraction and heart rate were preserved. In contrast, mice lacking the C3aR in leptin receptor–expressing neurons showed improved diastolic parameters, associated with lower heart weight and reduced myocardial fibrosis (Fig. 5a–i).

**Fig. 5.**
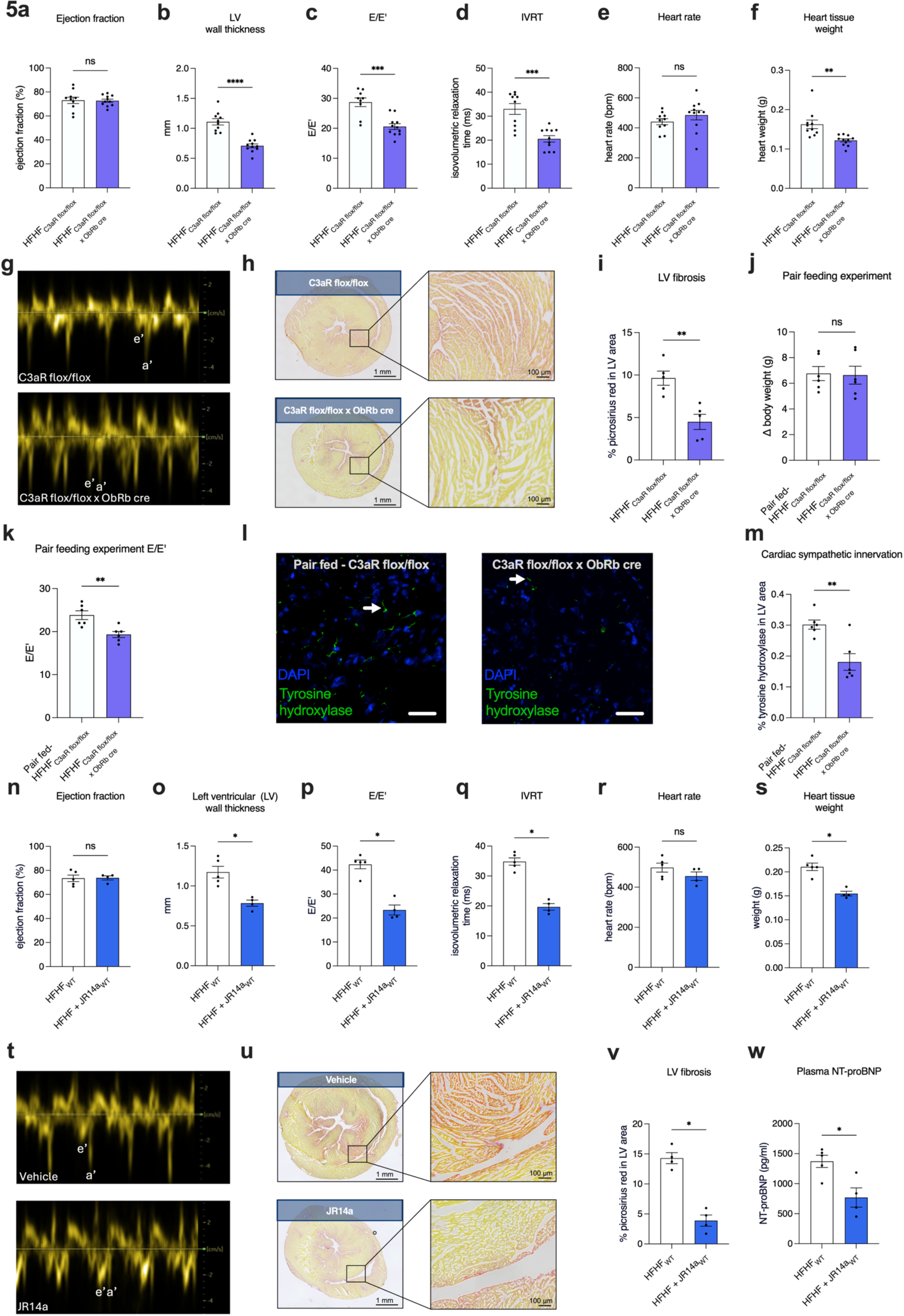
Neuronal C3aR deletion and pharmacological C3aR targeting attenuate obesity-associated diastolic dysfunction and myocardial remodeling. **a–f**, Echocardiographic assessment of cardiac function and remodeling in C3aR^flox/flox^ × ObRb-Cre mice and C3aR^flox/flox^ littermate controls after 16 weeks of high-fat/high-fructose (HFHF) diet. Shown are ejection fraction (a), left ventricular (LV) wall thickness (b), E/E′ (c), isovolumetric relaxation time (IVRT) (d), heart rate (e), and heart weight (f). g, Representative tissue Doppler images illustrating e′ and a′ waves in the indicated genotypes. h, Representative picrosirius red staining of LV sections (overview and higher magnification) from HFHF-fed C3aR^flox/flox^ controls and C3aR^flox/flox^ × ObRb-Cre mice. i, Quantification of LV fibrosis as % picrosirius red–positive area in LV. j–k, Pair-feeding experiment to separate body-mass-dependent effects from additional mechanisms. C3aR^flox/flox^ control mice were pair-fed (calorie-restricted) to match body weight gain of HFHF-fed C3aR^flox/flox^ × ObRb-Cre mice. Shown are Δ body weight (**j**) and E/E′ (**k**). **l,** Representative tyrosine hydroxylase immunostaining of LV sections from the pair-feeding cohort. White arrows point toward sympathetic nerves. Scale bar: 25 µm. **m,** Quantification of cardiac tyrosine hydroxylase-positive area, indicating sympathetic innervation. **n–s,** Therapeutic C3aR targeting with JR14a in WT mice suffering from HFHF diet-induced obesity. WT mice were maintained on HFHF for 18 weeks and then treated with vehicle or JR14a for 12 weeks while continuing HFHF. Shown are ejection fraction (**n**), LV wall thickness (**o**), E/E′ (**p**), IVRT (**q**), heart rate (**r**), and heart weight (**s**) at week 30. **t,** Representative tissue Doppler images (e′ and a′ waves) from vehicle- and JR14a-treated WT mice. **u,** Representative picrosirius red staining of LV sections from vehicle- and JR14a-treated WT mice. **v,** Quantification of LV fibrosis as percent picrosirius red–positive area in LV. **w,** Plasma NT-proBNP levels at week 30. Data are presented as mean ± SEM. Sample sizes: **a–f**: n=10-11; **i**: n=5; **j–m**: n=6; **n–s**: n=4-5; **v-w**: n=4-5. Statistical differences between groups were assessed using unpaired Student’s t-test (**a-f**) or Mann–Whitney U-test (**i-k**, **m-s**, **v-w**), ns: not significant, *p<0.05, **p<0.01, ****p<0.0001.

To separate weight-dependent from weight-independent effects, we pair-fed control mice to match the reduced food intake and body-weight gain of neuronal C3aR-deficient animals. Despite similar body weight under pair-feeding conditions, knockout mice maintained lower E/E′ values, supporting a cardiac protective mechanism beyond reduced weight gain alone (Fig. 5j-k). Cardiac tyrosine hydroxylase staining was reduced in pair-fed neuronal C3aR-deficient mice, consistent with altered sympathetic innervation (Fig. 5l–m), which is in line with prior work linking hypothalamic leptin signalling to sympathetic cardiovascular output^7,29^.

Importantly, we observed a similar protective effect in response to therapeutic pharmacological C3aR targeting in obese WT mice, which attenuated diastolic impairment, fibrosis and NT-proBNP elevation (Fig. 5n-w). Together, these data place neuronal C3aR signalling at the intersection of hypothalamic inflammation, leptin resistance and obesity-associated cardiac dysfunction.

### Obesity-associated complement activation in humans

To further place our findings into a translational context, we examined complement profiles in human obesity. In a large real-world cohort, obese individuals exhibited higher circulating C3, C4 concentrations and showed higher complement activation, indicated by C3c, than lean controls, consistent with prior clinical observations^14,30,31^ (Extended Data Fig. 5a-c and Supp. Table 2). In an independent longitudinal bariatric cohort, C3c declined after surgery-associated weight loss (Extended Data Fig. 5d and Supp. Table 3), consistent with systemic adiposity being associated with higher circulating complement activation markers that decrease with weight loss.

## Discussion

In this study, we implicate neuronal C3aR signalling in hypothalamic ObRb-expressing neurons as a previously unrecognized, important contributor to leptin resistance, diet-induced obesity and downstream cardiometabolic remodelling (Extended Data Fig. 6). Nutritional excess was associated with local hypothalamic complement activation, and C3a directly engaged neuronal C3aR to induce SOCS3 and attenuate leptin receptor signalling. Genetic deletion of C3aR in ObRb-expressing neurons preserved leptin responsiveness, reduced hyperphagia, protected against diet-induced obesity and improved glucose homeostasis. Pharmacological targeting of C3aR prevented obesity development and restored leptin sensitivity in established disease. Beyond metabolism, neuronal C3aR signalling also promoted obesity-associated diastolic dysfunction, myocardial fibrosis and cardiac sympathetic innervation, supporting a brain–heart mechanism of cardiac protection that extends beyond mere weight reduction.

Our findings extend current models of hypothalamic leptin resistance, which include impaired blood–brain barrier transport, receptor downregulation, lipotoxicity, endoplasmic reticulum stress and inflammatory signalling cascades such as IKK–NF-κB^4,5,32–35^. A common endpoint of these mechanisms is the induction of negative regulators including SOCS3 and PTP1B ^36–38^. Our data add complement-derived C3a as a distinct upstream signal, inducing SOCS3 through neuronal C3aR activation. This pathway integrates innate immune activation into the JAK– STAT–SOCS3 feedback circuit that controls leptin responsiveness. Thus, neuronal C3aR signalling acts as a neuroimmune amplifier of leptin resistance rather than a simple downstream correlate of hypothalamic inflammation.

A conceptual advance of the study is our observation that complement is activated locally within hypothalamic feeding circuits during obesogenic stress. C3 cleavage generates soluble C3a and activated C3 fragments detected in the ARC. Our focus on the C3a/C3aR axis was motivated by receptor-mediated neuronal signalling. Glial profiling suggests that both microglia and astrocytes contribute to hypothalamic C3 production. However, obesity preferentially increases astrocytic C3. Inflammatory cues also increase neuronal C3aR surface expression, consistent with a feed-forward circuit, in which nutritional excess promotes gliosis and complement activation, while simultaneously sensitizing hypothalamic neurons to C3a. Together these observations support a paracrine model in which locally generated C3a acts on leptin-sensitive neurons to induce SOCS3 and impair leptin signalling.

Our ObRb-Cre–based genetic model also helps to reconcile prior studies using global C3aR deficiency. Schadt et al. reported altered metabolic phenotypes in global C3aR-deficient mice, even under standard diet conditions^12^, whereas the tdTomato-C3aR^flox/flox^ x ObRb-Cre model studied here showed no body-weight phenotype under chow. Conversely, Mamane et al. observed only transient protection from diet-induced obesity in global C3aR-deficient mice^13^, whereas neuron-specific C3aR deletion resulted in sustained protection throughout HFHF exposure. These differences likely reflect distinct genetic strategies: global C3aR deficiency perturbs complement signalling in immune and peripheral metabolic compartments. In contrast, the ObRb-directed deletion used here points to a leptin-responsive neuronal component that becomes especially relevant under nutritional excess.

Obesity and leptin resistance are increasingly linked to diastolic dysfunction and myocardial remodelling^28,39^. The strong cardiac phenotype that we observed in response to nutritional excess broadens the physiological relevance of neuronal complement signalling. As hypothalamic leptin signalling interferes with autonomic output, leptin-driven sympathetic activation provides a plausible route through which central metabolic dysfunction influences peripheral cardiovascular disease^29^. Within this framework, our results suggest that neuronal C3aR signalling is not only permissive for hyperphagia and obesity but also contributes to obesity-associated diastolic impairment and fibrosis. The pair-feeding experiments were particularly informative, dissociating body-mass effects from additional mechanisms, while reduced cardiac tyrosine hydroxylase staining is consistent with altered sympathetic innervation as one candidate mediator. Thus, our data support a neuroimmune brain–heart axis linking hypothalamic inflammation to cardiometabolic end-organ disease.

The extent to which high-fat feeding alone in C57BL/6J mice recapitulates diastolic dysfunction remains debated and depends on diet composition, duration and experimental conditions^40^. Notably, in the present study, animals were exposed to a combined high-fat/high-fructose dietary challenge for up to 30 weeks, which imposes a more pronounced cardiometabolic stress compared to high-fat feeding alone. Previous studies have shown that obesogenic diets enriched in fat and simple sugars can promote cardiac remodeling and diastolic dysfunction^41–44^. Consistent with a biologically meaningful cardiac phenotype, NT-proBNP levels were reduced by C3aR modulation, supporting the presence of modifiable cardiometabolic stress in this model. Accordingly, we interpret our cardiac findings conservatively as obesity-associated diastolic impairment and remodeling rather than a complete heart failure with preserved ejection fraction (HFpEF) phenotype.

The translational implications of the preclinical model are strengthened by the human data. In a large real-world cohort, individuals with obesity showed higher circulating C3, C4 and C3c, consistent with previous reports linking complement dysregulation to obesity, insulin resistance and metabolic liver disease^14,30,45,46^. In a longitudinal bariatric cohort, C3c declined after weight loss, supporting complement activation as a dynamic biomarker of adiposity and metabolic improvement. These systemic readouts do not directly report on brain complement activation, but they provide an important clinical context for complement dysregulation as a hallmark of obesity and support the plausibility of complement-targeted interventions in cardiometabolic disease. Future translational work integrating biomarker profiling with advanced imaging approaches to infer brain complement activation may help bridge this gap.

Recent incretin-based therapies have substantially improved the clinical management of obesity and obesity-associated heart failure with preserved ejection fraction^47,48^. However, despite their efficacy in promoting weight loss and improving cardiometabolic outcomes, important limitations remain, including gastrointestinal adverse effects, incomplete responses in a subset of patients and attenuation or loss of therapeutic benefit after treatment discontinuation^49^. These considerations highlight the need for complementary or mechanistically distinct interventions. In this context, neuronal C3aR signalling emerges as a candidate pathway that could complement existing incretin-based strategies, particularly in patients with persistent leptin resistance or cardiometabolic remodelling despite weight loss.

Targeting C3aR may also provide advantages over broader anti-inflammatory interventions. Inhibition of upstream innate immune pathways can compromise host defence^50^, whereas selective modulation of C3aR is expected to leave key complement effector functions such as C3b-mediated opsonization and terminal pathway activation intact. Nevertheless, C3aR is expressed beyond neurons, including in tissue-resident macrophages, endothelial cells and regenerative compartments^9,51–53^. Thus long-term systemic C3aR modulation will require careful safety evaluation. At this point, we cannot exclude that systemic pharmacological C3aR inhibition engages additional mechanisms outside the hypothalamus. The concordance between neuron-specific genetics and pharmacology supports the central conclusion, but future work with alternative chemotypes, CNS-penetrant or neuron-targeted delivery and direct measurements of sympathetic output will be important. C3aR can additionally be engaged by ligands such as TLQP-21 ^54^, underscoring that the receptor may integrate broader metabolic signals beyond canonical complement activation.

Several limitations should be acknowledged. First, while we establish neuronal C3aR expression in the hypothalamus and define its effects on leptin signalling, further work is needed to map signalling across specific leptin-responsive subpopulations, including AgRP and proopiomelanocortin (POMC) neurons. Second, our cardiac phenotyping focuses on echocardiographic diastolic indices, fibrosis and sympathetic innervation; additional haemodynamic studies could further refine alignment with clinical heart failure. While tyrosine hydroxylase staining provides information on structural sympathetic innervation, it does not establish functional autonomic tone, which will require direct physiological assessment (e.g. telemetry or norepinephrine turnover) in future studies. Moreover, complementary multi-hit models incorporating haemodynamic stressors, ageing or renal dysfunction may more fully recapitulate the complexity of human HFpEF. Third, systemic complement measurements in human cohorts provide translational context but do not establish central complement activation or causality. In addition, JR14a was originally developed as a C3aR antagonist and, in our mHypoE-N46 assay, prevented the C3a-mediated attenuation of leptin action. However, recent structural and functional studies indicate that JR14a can also activate C3aR and promote receptor internalization in other systems^55–57^. Thus, the in-vivo data should be interpreted as pharmacological C3aR modulation rather than definitive receptor blockade. The concordance with ObRb-directed C3aR deletion supports the receptor-level interpretation, but future studies with alternative chemotypes will be important. Finally, although our validation analyses did not provide evidence for relevant off-target recombination in the examined peripheral compartments, the ObRb-Cre strategy does not confer strict hypothalamic region restriction. Future studies using viral or circuit-specific approaches will help to further refine the anatomical and temporal specificity of C3aR signalling.

In conclusion, our data support neuronal C3aR signalling as a modifiable neuroimmune pathway that contributes to impaired hypothalamic leptin responsiveness, hyperphagia, obesity and obesity-associated cardiac remodelling. These findings nominate C3aR signalling as a candidate therapeutic axis for further evaluation in cardiometabolic disease, potentially alongside existing obesity therapies.

## Methods

### Animal Studies

All animal care and experimental procedures were performed in accordance with the NIH guidelines for the care and use of laboratory animals. The animal studies are reported according to the ARRIVE guidelines. All animal experiments were approved by the animal ethics committee of the local regulatory authority (Ministry of Agriculture, Rural Areas, European Affairs and Consumer Protection Schleswig-Holstein. Protocol # 44-9/25, 7-2/22).

B6.129(Cg)-Leprtm2(cre)Rck/J) (ObRb Cre) mice were obtained from Charles River Laboratories (Sulzfeld, Germany) and C.C3aRtm1JKo (C3aR tdTomato floxed) mice were generated as described before^19^. Before crossing with ObRb Cre mice, C3aR tdTomato floxed mice were backcrossed to C57BL/6J background for 10 generations. C57BL/6J mice were obtained by Charles River Laboratories (Sulzfeld, Germany). Males and females in approximately equal numbers were used for the experiments. Mice were housed 4–5 per cage in a pathogen-free mouse facility with ad libitum access to food and water on a 12-hour light/12-hour dark cycle. Animals were fed either a control diet (chow, ssniff E15755) or a high-fat diet containing 45% energy from fat, supplemented with 22% fructose (ssniff E15756), referred to as a high-fat/high-fructose (HFHF) diet. Mice entered indicated diet feeding at 7-8 weeks of age. Investigators performing echocardiographic, histological and image-based analyses were blinded to genotype/treatment during acquisition and quantification. Animals were allocated to treatment groups computer-based randomization. C57BL/6J mice shown in Fig. 4a-f and Fig. 4h-m were treated for the indicated time with JR14a (10 mg/kg/d) in drinking water ^21^. The concentration was adjusted based on continuous monitoring of water consumption to ensure accurate and consistent delivery of the compounds. At the end of the experiment, terminal blood samples (500 µl) were collected under anesthesia using ketamine (100 mg/kg, i.p.) and xylazine (10 mg/kg i.p.) via the retro-orbital venous plexus, and plasma was prepared for further analyses.

### Leptin resistance test

To evaluate leptin sensitivity, a leptin resistance test was performed at week 14 for Extended Data Fig. 1f and Fig. 3o as well as at week 18 for Fig 4g and week 28 for Fig. 4m. For that, body weight was measured over five consecutive days following repeated intraperitoneal (i.p.) injections of leptin (1 mg/kg body weight in 0.9% NaCl). The i.p. injections were administered twice daily at 17:00 and 23:00 from day 1 to day 5 of the experiment, as previously described^18^.

### Leptin-induced pSTAT3 activation in hypothalamic neurons

To identify leptin-responsive neurons in vivo, tdTomato-C3aR reporter mice were injected intraperitoneally with recombinant leptin (3 mg/kg body weight). After 30 minutes, mice were euthanized and brains were collected, fixed in 4% paraformaldehyde and processed for immunofluorescence staining.

### Glucose tolerance test

The glucose tolerance test (GTT) was performed at week 12 for Extended Data Fig. 1c, Fig. 3k-l and at week 8 for Fig. 4e as well as at week 25 for Fig. 4k. Therefore, food was withdrawn 12 hours prior to the procedure, while water remained available ad libitum. Following a baseline blood sample from the lateral tail vein, a glucose solution (1.5 g/kg body weight in 0.9% NaCl) was administered via intraperitoneal injection. Blood glucose levels were measured at 20, 40, 60, 90, and 120 minutes after glucose administration using an Aviva Accu Chek device. Approximately 10 µl of blood was collected at each time point. For blood collection, animals were briefly restrained in a mouse injection holder. To dilate the tail vein, the tail was immersed in warm water (maximum 45 °C) for a few seconds, sparing the sensitive tail tip. Area of the curve (AOC) for the GTT in Extended Data Fig. 1c, Fig. 3k, Fig. 4e and Fig. 4k was calculated by subtracting the baseline glucose value from each time point prior to integration, as described previously ^58^. This method accounts for differences in fasting glucose levels and provides a more accurate assessment of glucose tolerance than traditional area under the curve (AUC) analysis.

### Insulin tolerance test

The insulin tolerance test (ITT) was performed at week 13 for Extended Data Fig. 1d, Fig. 3m-n and at week 9 for Fig. 4f as well as at week 26 for Fig. 4l. Therefore, food was withdrawn 5.5 hours prior to the experiment, while water remained available ad libitum. Following a baseline blood sample, mice received an injection of insulin (1 IU/kg body weight, equivalent to 2 µl/g body weight of insulin solution; Insuman, Aventis) in 100 µl of 0.9% NaCl via intraperitoneal injection. Blood glucose levels were measured at 15, 30, 60, and 90 minutes after injection. At each time point, approximately 10 µl of blood was collected by incision of the lateral tail vein. For safe blood collection, animals were briefly restrained in a mouse injection holder. To dilate the tail vein, the tail was briefly immersed in warm water (maximum 45 °C), sparing the sensitive tip. AOC for the ITT in Extended Data Fig. 1d, Fig. 3n, Fig. 4f and Fig. 4l was calculated by subtracting the baseline glucose value from each time point prior to integration, as described previously ^58^.

### Indirect calorimetry

Indirect calorimetry was used at week 10 to assess metabolic rate by measuring oxygen consumption (VO₂) and carbon dioxide production (VCO₂) in individually housed mice. Animals were placed in specialized metabolic chambers (TSE Systems), with continuous measurement of incoming and outgoing O₂ and CO₂ concentrations. From these values, the respiratory exchange ratio (RER) was calculated as the ratio of VCO₂ to VO₂ over a defined time period ^59^. Additional parameters including cage temperature, food and water intake were continuously recorded. Locomotor activity was monitored using an infrared beam-based system (InfraMot®, TSE Systems). For data analysis, raw CSV output files from the calorimetry system were aggregated and time-aligned across all treatment groups. Daytime (6:00–18:00) and nighttime (18:00–06:00) periods were defined consistently and marked across datasets. RER, energy expenditure, energy intake and locomotion were analyzed over a 48 h post-acclimatization period, RER, locomotion and energy expenditure were averaged per animal for daytime and nighttime intervals over two days. Total energy intake (kcal) was computed from food weight using the caloric density of the HFHF diet (4,920 kcal/kg).

### Pair-feeding experiments

For pair-feeding experiments, C3aR^flox/flox^ control mice were single-housed and received the same HFHF diet as C3aR^flox/flox^ × ObRb-Cre mice. Food intake of the C3aR^flox/flox^ × ObRb-Cre group was recorded over consecutive 24h intervals, and C3aR^flox/flox^ control mice were provided the corresponding amount of HFHF diet for the subsequent 24 h. Food was provided once daily in the afternoon immediately before the onset of the dark phase. Intake was determined from food disappearance over the preceding 24 h (food provided minus residual food and visible spillage). Water remained available ad libitum throughout the experiment. Body weight was monitored longitudinally during the pair-feeding period.

### Echocardiography

Transthoracic echocardiography was performed at predefined time points to assess cardiac function with emphasis on diastolic dysfunction. Mice were anesthetized with an intraperitoneal ketamine injection (100 mg/kg body weight in 0.9% NaCl), the anterior chest was depilated using a chemical hair remover, and animals were placed on a temperature-controlled heating pad. Heart rate and electrocardiogram were continuously monitored. Echocardiography was performed using a GE i13L transducer on a GE Vivid E9 system. Parasternal long-and short-axis views were used to assess left ventricular (LV) structure and systolic function, including LV wall thickness and ejection fraction. Diastolic function was evaluated in the apical four-chamber view. Early diastolic mitral annular velocity was measured by tissue Doppler imaging at the septal mitral annulus to calculate E/E′. Isovolumetric relaxation time (IVRT) was determined as the interval between aortic valve closure and the onset of mitral inflow.

### Culture of Embryonic Mouse Hypothalamic Neurons (mHypoE-N46)

Immortalized murine hypothalamic neurons (mHypoE-N46) were purchased from Biozol (Biozol, CED-CLU138). Cells were cultured in proliferation medium consisting Dulbecco’s Modified Eagle Medium (DMEM) (ThermoFisher, 11965092) with 10% fetal bovine serum (FBS) and 1% penicillin-streptomycin (Thermo Fisher Scientific, 15140122). Cells were maintained at 37°C in a humidified atmosphere containing 5% CO₂. The culture medium was refreshed every 3-4 days, and cells were passaged upon reaching 90% confluence.

### qPCR Analysis

#### Murine hypothalamus

Mice were euthanized and transcardially perfused with 10 ml ice-cold PBS. Brains were removed, and hypothalami rapidly dissected and homogenized in TRIzol reagent (Sigma-Aldrich) for 5min at room temperature (RT). Phase separation was induced by chloroform incubation for 2min at RT, followed by centrifugation for 15min at 11,600x g at 4°C. After collecting the upper aqueous phase, RNA was precipitated with ethanol. Total RNA was purified using the RNeasy Mini Kit (Qiagen) including on-column DNase I digestion to eliminate genomic DNA contamination, according to the manufacturer’s instructions. RNA concentration and purity were determined using a NanoDrop spectrophotometer.

Complementary DNA (cDNA) was synthesized using the High-Capacity cDNA Reverse Transcription Kit (Thermo Scientific, 4368814) following the manufacturer’s protocol. To prevent RNA degradation, RiboLock RNase Inhibitor (Thermo Scientific) was included during cDNA synthesis. A total of 10 ng of cDNA was used for quantitative analysis. Quantitative PCR was performed in 384-well plates on a Applied Biosystems™ 7900HT. Expression of SOCS3 was quantified using gene-specific primers (forward: 5′-ACC AGC GCC ACT TCT TCA CG-3′; reverse: 5′-GTG GAG CAT CAT ACT GAT CC-3′). Gene expression levels were normalised using the formula 2-ΔCT using the gene expression levels of murine y-Actin as housekeeping gene.

#### Analysis of C3aR expression in peripheral tissues

To assess whether ObRb-Cre–mediated recombination causes unintended C3aR deletion in peripheral metabolic tissues and heart tissue, we performed quantitative PCR analysis on perigonadal white adipose tissue, general white adipose tissue, brown adipose tissue, pancreas, liver, and whole heart samples from Cre-positive and Cre-negative C3aR-tdTomato floxed mice. Following transcardial perfusion with PBS, tissues were rapidly dissected, snap-frozen in liquid nitrogen, and cryo-ground to a fine powder. The powder was lysed with QIAzol™ (QIAGEN) and disrupted using a cannula attached to a 1 mL syringe. Chloroform was added, followed by shaking and a brief incubation of 5 min at RT. The mixture was centrifuged at 15,000 × g for 15 minutes at 4°C to separate the aqueous and organic phases. Ethanol was then added to the aqueous phase, and the solution was resuspended and transferred to a RNeasy Mini Kit (Qiagen) column, followed by on-column DNase I digestion. RNA concentration and quality were assessed using a Nanodrop spectrophotometer and Qubit 4 fluorometer (Thermo Scientific). cDNA was synthesized using the High-Capacity cDNA Reverse Transcription Kit (Thermo Scientific, 4368814) according to the manufacturer’s protocol. A total of 20 ng of cDNA was used for quantitative analysis. *C3ar1* mRNA levels were quantified using a gene expression master mix (Thermo Scientific, 4369514) and specific assays for *C3ar1* (Mm01184110_m1) and *Cdkn1b* (Mm00438167_g1). Quantification was performed on a QuantStudio 7 Flex (Applied Biosystems).

#### Analysis of SOCS3 expression in mHypoE-N46 cells

To evaluate SOCS3 expression upon C3aR activation, mHypoE-N46 cells were stimulated for 4 hours in proliferation medium either alone or supplemented with 100 nM C3a, or with 5 nM interleukin-6 (IL-6). Following stimulation, cells were washed with PBS and lysed using RNeasy Lysis Buffer (Qiagen, 79216) supplemented with 1:100 β-mercaptoethanol. The cells were then homogenized and transferred to a RNeasy Mini Kit (Qiagen) column. On-column DNase I digestion followed. The RNA concentration and quality were then assessed using a Nanodrop spectrophotometer and Qubit 4 fluorometer (Thermo Scientific), after which cDNA was synthesized using a High-Capacity cDNA Reverse Transcription Kit (Thermo Scientific, 4368814) according to the manufacturer’s protocol. Gene expression analysis was performed using a gene expression master mix (Thermo Scientific, 4369514) and specific assays for *PTP1B* (Mm00448427_m1), and *SOCS3* (Mm00545913_s1). Quantification was performed on a QuantStudio 7 Flex (Applied Biosystems). A total of 20 ng of cDNA was used for quantitative analysis. Gene expression levels were normalized using the formula 2^-ΔCT using the gene expression levels of murine *B2m* (Mm00437762_m1) as reference gene.

### Flow Cytometric Analysis

#### C3aR expression in mHypoE-N46 cells

To assess C3aR expression in murine hypothalamic neurons, mHypoE-N46 cells were incubated in proliferation medium with or without 5nM IL-6 for 12h. Cells were detached using 5mM EDTA in phosphate-buffered saline (PBS) for 5 min at RT and washed with ice-cold PBS containing 1% bovine serum albumin (BSA). Cell suspensions were centrifuged at 300xg for 5min at 4°C. For surface staining, cells were incubated with an unconjugated primary anti-C3aR antibody (1:100, HycultBiotech, HM1123) in PBS containing 1% BSA and 1% normal donkey serum (NDS) for 45min at 4°C in the dark. After washing, cells were fixed with 4% paraformaldehyde for 15min at RT and washed twice. Intracellular staining was performed by incubating cells with anti-C3aR (1:100, HycultBiotech, HM1123) and anti-beta-III tubulin-Alexa Fluor 488 (1:100, ThermoFisher, 53-4510-82) in permeabilization buffer consisting of BD Perm/wash buffer (1:10 in PBS, BD Bioscience, 554723), 1% BSA and 1% NDS for 1h at 4°C. After two washes, cells were incubated with donkey anti-rat IgG–Alexa Fluor 594 secondary antibody (1:1000, ThermoFisher, A-21209) in permeabilization buffer for 1h at 4°C. Cells were then washed twice, resuspended in PBS containing 1% BSA, and stained with DAPI (1:3000, ThermoFisher, D1306) for 1 min prior analysis. Data were acquired on CytoFLEX S flow cytometer (Beckman Coulter Life Sciences) and analyzed using the Cytobank platform (Beckman Coulter Life Sciences).

#### Analysis of microglia and astrocytes

To quantify microglial and astrocytic and assess local complement C3 production *in-vivo*, C57BL/6J mice were euthanized and transcardially perfused with 10 ml ice-cold PBS. Brains were removed, and the hypothalamus dissected, minced into small pieces in ice-cold PBS, and digested with Dispase/Collagenase (1 mg/ml; Sigma Aldrich) and DNase I (1 mg/ml, Sigma Aldrich) at 37°C for 50min on a thermoshaker at 800 rpm. Following gentle mechanical dissociation, cell suspensions were filtered through a 70-µm cell strainer, washed with FACS Buffer consisting of PBS supplemented with BSA at a 1:200 dilution, and centrifuged at 300xg for 10min at RT. Supernatants were discarded. Cells were resuspended in 30% Percoll Solution, under-layered with 70% Percoll Solution and centrifuged at 340 × g for 20min at RT without brake. The interphase was collected, washed with Post-Percoll Solution containing HBSS supplemented with 5% FBS and 10 µM HEPES, and centrifuged at 300 × g for 10min at 4°C.

Cells were resuspended in RBC lysis buffer (1:10 in H2O; BioLegend) for 5min, washed and resuspended in FACS buffer. For surface staining, cells were blocked with FcgR -block (1:100; Miltenyi Biotec) on ice for 10min, followed by Zombie viability dye (1:1000; BioLegend, 423105) for 10min at 4°C in the dark. Cells were incubated with anti-CD68 (1:50, BioLegend,137008), anti-CD45 (1:50, BioLegend, 103140), anti-CD11b (1:50; BioLegend, 101208), anti-C3aR (1:50; Santa Cruz Biotechnology, SC-133172 FITC) antibody cocktail and Hoechst dye (1:1000; Sigma-Aldrich, 63493) for 30min at 4°C in the dark.

After washing, cells were fixed with 1% PFA in FACS Buffer for 20min at RT in the dark. Following two washes, cells were incubated with anti-C3 (1:50; Novus Biologicals, NB200-540PCP) and anti-GFAP Alexa-Fluor 568 (1:50; Abcam, ab201736) antibodies in flow cytometry permeabilization buffer (R&D Systems, FC005) for 30min at RT, protected from light. Cells were washed with flow cytometry permeabilization buffer, centrifuged for 5min at 300xg, resuspended in FACS Buffer, and analyzed by flow cytometry.

#### Analysis of C3aR expression in bone marrow and peripheral blood

To determine whether ObRb-Cre–mediated recombination induces off-target C3aR deletion outside the hypothalamus, we assessed C3aR expression in bone marrow-derived mesenchymal stromal cells (MSCs), total bone marrow leukocytes, peripheral neutrophils, and macrophages from both, Cre-positive and Cre-negative C3aR-tdTomato floxed mice. Bone marrow cells were isolated by flushing femur and tibia shafts with ice-cold FACS buffer containing 2 mM EDTA using a 25G needle and a 12 cc syringe. Remaining bone fragments were minced and filtered through a 70 µm cell strainer. Cells were pelleted by centrifugation at 500 × g for 7 min at 4 °C, erythrocytes were lysed with RBC lysis buffer (BioLegend, 420301) for 5 min, and remaining cells were washed twice in FACS buffer. For surface staining, cells were incubated with FcgR block (1:10, Miltenyi Biotec, 130-092-575) on ice for 10 min, followed by staining with a pre-mixed antibody cocktail for CD11b (BioLegend, 101205), CD45 (BioLegend, 103138), Ly6C (BioLegend, 128049), Ly6G (Miltenyi Biotec, 130-128-232), F4/80 (BioLegend, 123131), Sca-1 (BioLegend, 108125), CD31 (BioLegend, 160214), allowing identification of MSCs (CD45⁻CD31⁻CD11b⁻Ly6G⁻F4/80⁻Sca-1⁺), for 30 min at 4 °C protected from light. After washing, cells were resuspended in FACS buffer and analyzed by flow cytometry.

For peripheral blood analysis, fresh blood was collected into ACD-containing tubes. After FcgR block (1:10, Miltenyi Biotec) for 10 min at RT, cells were stained with by Zombie viability dye (1:1000, BioLegend, 423105) for 10 min, followed by incubation with the antibody cocktail (CD11b (BioLegend, 101205), CD45 (BioLegend, 103138), Ly6G (Miltenyi Biotec, 130-128-232), F4/80 (BioLegend, 123131), for 30 min at 20 °C. Red blood cells were lysed with RBC lysis/fixation solution (BioLegend, 422401) for 15 min in the dark at RT, then cells were washed, resuspended in FACS buffer, and analyzed by flow cytometry. Data were analyzed to compare C3aR-tdTomato expression between Cre-positive and Cre-negative animals to confirm tissue specificity.

### Immunofluorescence Staining

#### Immunofluorescence staining of C3aR in mHypoE-N46 cells

Expression levels of C3aR and leptin receptor in murine hypothalamic neurons were assessed by immunofluorescence staining. Cells were washed three times with PBS and fixed with BD Cytofix/Cytoperm buffer (BD Bioscience, 554722) for 15min at RT. After two washes with blocking/permeabilization buffer consisting of BD Perm/wash buffer (1:10 in PBS, BD Bioscience, 554723) and 1% BSA, cells were incubated overnight at 4°C with an unconjugated primary antibody against C3aR (1:100, HycultBiotech, HM1123) or isotype control (1:100, Thermo-Fisher, 02-9688) and a conjugated primary antibody against leptin receptor (1:100, Santa Cruz Biotechnology, SC-8391) diluted in blocking/permeabilization buffer containing 1% NDS. Following three washes, cells were incubated with donkey anti-rat IgG–Alexa Fluor 488 secondary antibody (1:1000, ThermoFisher, A-21208) in block/perm buffer supplemented with 1% NDS for 1 hour at RT. Cells were washed three times and stained with DAPI (1:3000, ThermoFisher, D1306) for 1min. Images were acquired using a Keyence BZ-X810 fluorescence microscope.

#### Immunofluorescence staining of murine hypothalamus

Following transcardiac perfusion with PBS and fixation in 4% paraformaldehyde (PFA), brains were dissected, cryopreserved, and coronally sectioned at 14 µm thickness using a cryostat (Leica, CM30503). Sections were mounted on adhesion microscopy slides (Epredia, J1800AMNZ) and stored at –20 °C until use. Prior to staining, sections were air-dried for 30 min at RT, rehydrated in PBS for 30 min and washed three times in PBS for 10 min each. To block nonspecific binding, sections were incubated in 5% normal goat serum (Cell Signaling, 5425) in PBS-T (0.5% Triton X-100, 0.1% Tween-20) for 1h at RT. Unconjugated primary antibodies including rat anti-GFAP (Invitrogen, 13-0300), rabbit anti-Iba1 (FUJIFILM Wako, 019-19741) and mouse anti-MAP2 (Merck, MAB3418) were diluted 1:500 in PBS-T containing 1.5% normal goat serum and incubated overnight at 4 °C. After five washes in PBS, sections were incubated with species-specific Alexa Fluor-conjugated secondary antibodies including goat anti-mouse Alexa Fluor 647 (Invitrogen, A-21236), goat anti-rat Alexa Fluor 488 (Invitrogen, A-11006) and goat anti-rabbit Alexa Fluor 647 (Abcam, ab150079) diluted 1:1000 in PBS-T with 1.5% normal goat serum for 2h at RT.

In parallel, activated fragments of the mouse complement protein C3 were detected using an anti-C3b/iC3b/C3c antibody (Hycult, HM1065). Sections were counterstained with DAPI (Carl Roth GmbH, 6335.1) and mounted with fluorescence mounting medium (Dako, Agilent, S3023). For phosphorylated STAT3 (pSTAT3) detection as a readout of leptin receptor activation and NeuN to identify neurons phospho-Stat3 (Tyr705) rabbit antibody (Cell Signaling Technology, 9145S) and anti-NeuN antibody, clone A60 (Merck, MAB377X) were used. Co-localization of pSTAT3 with tdTomato-labelled C3aR was analyzed in the hypothalamus to assess C3aR expression in leptin-responsive neuronal populations. All slides were dried and imaged using Keyence BZ-X810 fluorescence microscope or Leica Stellaris 8 confocal microscope.

#### Immunofluorescence staining of murine heart tissue

Following transcardiac perfusion with PBS, hearts were collected, cryopreserved, and sectioned into 5 µm slices using a cryotome at –20 °C. Sections were placed on glass slides for immunofluorescence analysis of catecholaminergic innervation. Tissue sections were washed three times in PBS for 10 minutes each and blocked for 1 hour at room temperature in PBS containing 10% normal goat serum (NGS) and 0.3% Triton-X. Sections were incubated overnight at 4 °C with anti-tyrosine hydroxylase (TH) primary antibody (Abcam, Ab152) 1:500 in PBS containing 1% NGS and 0.3% Triton-X, followed by three washes in PBS for 10 minutes each. Sections were then incubated for 2 hours at room temperature with goat anti-rabbit Alexa Fluor 488 secondary antibody and DAPI 1:3000 in PBS containing 1% NGS and 0.3% Triton-X, followed by three final washes in PBS. Sections were mounted in fluorescence mounting medium (Dako, Agilent, S3023) and imaged using a Leica Stellaris 8 confocal microscope.

### Histochemical staining

#### Picro-Sirius Red Staining

Following transcardiac perfusion with PBS, hearts were collected, cryopreserved, and sectioned into 5 µm thick slices using a cryotome at −20°C. Sections were placed on glass slides for histological analysis of fibrosis. Picro-Sirius Red Staining was performed using the Picro-Sirius Red Stain Kit (Abcam, ab245887). Tissue sections were fully covered with 0.2% phosphomolybdic acid solution for 5 min, rinsed once in distilled water, and then stained with Picro-Sirius Red solution for 75 min at room temperature. Slides were quickly rinsed in two changes of 0.5% acetic acid, dehydrated sequentially in 70% ethanol, absolute ethanol, and cleared in xylene, before mounting with a xylene-based medium. Stained sections were imaged using a BZ-X810 microscope (Keyence).

### Enzyme-linked Immunosorbent Assays

#### Evaluation of murine plasma leptin levels

Circulating leptin levels were assessed in plasma samples obtained from C57BL/6J obese and normal-weight mice. Plasma was separated by centrifugation at 10,000 × g for 4 minutes at 4°C. Leptin concentrations were measured using a mouse Leptin ELISA kit (R&D Systems, MOB00B) following the manufacturer’s protocol.

#### Evaluation of murine plasma NT-proBNP levels

Circulating NT-proBNP levels were quantified in plasma samples obtained from C57BL/6J mice receiving either vehicle or the C3aR modulator JR14a in combination with HFHF diet. Plasma was separated from whole blood by centrifugation at 10,000 × g for 4 minutes at 4°C. NT-proBNP concentrations were measured using a mouse NT-proBNP ELISA kit (Amsbio, AMS.E-EL-M0834-96T) according to the manufacturer’s instructions.

#### Complement C3a expression in hypothalami of WT-mice

For the analysis of C3a expression, brains were harvested from C57BL/6J obese and normal weight mice after transcardial PBS perfusion, and the hypothalamus was isolated. The hypothalamic tissue was lysed in 10% RIPA lysis buffer containing 1% Halt Protease and Phosphatase Inhibitor (ThermoFisher, 78440), and C3a levels were measured using a mouse complement C3a ELISA kit (Novus Biologicals, NBP2-70037) according to the manufacturer’s instructions. C3a concentrations were normalized to total protein content determined by a bicinchoninic acid (BCA) Protein Assay Kit (ThermoFisher, 23225).

#### Quantification of AgRP levels in mHypoE-N46 cells

Agouti-related peptide (AgRP) levels were quantified in cell lysates using a mouse AgRP ELISA kit according to the manufacturer’s protocol (ABclonal, RK02578). mHypoE-N46 cells were treated for 12h with either proliferation medium or glucose-free DMEM (ThermoFisher, ref: A1443001) supplemented with leptin (10nM; R&D Systems, reconstituted in 20 mM Tris-HCI, pH 8.0), leptin (10nM) and C3a (100nM; R&D Systems, reconstituted in PBS), or leptin (10nM), C3a (100nM), and the C3aR modulator JR14a (10 µM; MedChemExpress, HY-138161, dissolved in DMSO). Following treatment, the culture medium was carefully removed. Cells were lysed directly in the wells using RIPA buffer (Abcam, ab156034) supplemented with protease and phosphatase inhibitors (ThermoFisher, 78440) and incubated for 1h at 4°C at 800rpm. Lysates were centrifuged at 14,000 g for 10 min at 4 °C to remove cellular debris. Total protein concentrations were determined using a BCA Protein Assay Kit. AgRP levels were normalized to total protein concentration and presented relative to the glucose-free DMEM condition, which served as control.

### Retrospective cohort: complement profiling in obesity

We analyzed real-world data from the TriNetX research platform, a global federated network of anonymized electronic health records covering data from 2011 to 2024. 559,917 adult patients were retrospectively identified if plasma levels of complement components C3, C4, and C3c had been measured during routine clinical care. Both male and female patients were included, and sex was considered as a demographic variable in the dataset; no sex-based exclusion criteria were applied. Patients were categorized based on their body mass index (BMI) into an obesity group (BMI ≥30 kg/m²) and a lean control group (BMI within the normal range, 18.5–24.9 kg/m²). Demographic and clinical characteristics were extracted and are summarized in Suppl. Table 2. As TriNetX provides only de-identified data, no additional ethical approval was required for this analysis.

### Human bariatric surgery cohort

Plasma samples from obese patients undergoing bariatric surgery (gastric stapling) were collected before and 12 months after surgery. Peripheral blood samples were taken from 102 obese patients undergoing bariatric surgery. Both male and female patients were included. Blood samples were collected at prior to surgery (baseline) and approximately 12 months after surgery. All participants provided written informed consent. At baseline, 89 blood samples could be analyzed, at 12 months, 56 patient blood samples were still available for analysis. Plasma samples were collected after venipuncture into EDTA tubes, centrifuged at 1.500 × g for 15 min at 4 °C, and stored at –80 °C until analysis.

#### Plasma C3c analysis in bariatric surgery patients

Plasma C3c concentrations were quantified using an enzyme-linked immunosorbent assay (ELISA) kit (Hycult Biotech, Cat. No. HK368) following the manufacturer’s instructions. Briefly, samples were thawed, brought to room temperature, and diluted 1:50 in the kit’s dilution buffer. 100 µL of standards, controls, and diluted samples were incubated in antibody-coated micro-titer wells for 60 min at 20–25 °C and signal was detected using biotinylated tracer antibody and streptavidin–peroxidase conjugate. Absorbance was detected at 450 nm using an automated plate reader.

### Study approval

The study protocol for the human bariatric surgery cohort was approved by the institutional review board (Ethikkommission der Medizinischen Fakultät der Universität Heidelberg/ Mannheim, AZ 2021-602) before study commencement and conducted in accordance with the Declaration of Helsinki.

### Statistical analysis

Data were analyzed using GraphPad Prism 10 (GraphPad Software) or SPSS 25 (IBM). Sample sizes for animal experiments were planned based on a power analysis by G*Power using body weight gain as primary outcome parameter. Significance was considered when p < 0.05. Depending on the dataset and experimental design, different statistical methods were used as indicated in the Figure legends. Parametric statistics (e.g., t-Test, ANOVA) were only applied if assumptions were met, i.e., datasets were examined for Gaussian distribution by D’Agostino-Pearson test, aided by visual inspection of the data, and homogeneity of variances by Brown-Forsythe, Levene’s or F-test (depending on the statistical method used). If assumptions for parametric procedures were not met or could not be reliably assumed due to small sample size, non-parametric methods were used as indicated. Two-tailed tests were applied if not indicated otherwise. For multiple comparisons following ANOVA, Bonferroni-adjusted post hoc tests were applied. Energy expenditure was additionally analysed by ANCOVA with body mass as covariate; group × mass interaction was tested before interpreting the main effect of group.

## Data availability

Data supporting the findings of this study are available from the corresponding authors upon reasonable request.

## Acknowledgements

We thank Anke Constantz, Alexander Haschke, Maren Behrensen, Xinyu Hu and Rohit Chadhari for their excellent technical assistance, and Jens Mittag for his support with the TSE System.

## Author Contributions

Conceptualization: ER, CG, JK

MethodologyER, CG, ZA, WR, TZ, HL, JK, IE

Formal Analysis: ER, CG, MS, TR, KK

Investigation (conducting experiments, acquiring data): ER, CG, MS, TR, KK, VP, SMR, SG, JH, NLA, NA, JDE

Data Curation: ER, CG, MS, TR, NA, HL

Resources (reagents, materials, data access): AT, ZA, WR, MO, TZ, HL, HO, SGM, JK

Visualization: ER, CG

Writing – Original Draft: ER

Writing – Review & Editing: All authors

Project Administration: ER

Supervision: IE, JK

During the preparation of this work the authors used DeepL Pro in order to improve language and readability. After using this tool, the authors reviewed and edited the content as needed and take full responsibility for the content of the publication.

## References

1 Valenzuela, P. L. et al. Obesity and the risk of cardiometabolic diseases. Nat Rev Cardiol 20, 475–494, doi:10.1038/s41569-023-00847-5 (2023).

2 Jais, A. & Bruning, J. C. Hypothalamic inflammation in obesity and metabolic disease. J Clin Invest 127, 24–32, doi:10.1172/JCI88878 (2017).

3 Friedman, J. M. Leptin and the endocrine control of energy balance. Nat Metab 1, 754–764, doi:10.1038/s42255-019-0095-y (2019).

4 Thaler, J. P. et al. Obesity is associated with hypothalamic injury in rodents and humans. J Clin Invest 122, 153–162, doi:10.1172/jci59660 (2012).

5 de Git, K. C. & Adan, R. A. Leptin resistance in diet-induced obesity: the role of hypothalamic inflammation. Obes Rev 16, 207–224, doi:10.1111/obr.12243 (2015).

6 Bjorbaek, C., Elmquist, J. K., Frantz, J. D., Shoelson, S. E. & Flier, J. S. Identification of SOCS-3 as a potential mediator of central leptin resistance. Mol Cell 1, 619–625, doi:10.1016/s1097-2765(00)80062-3 (1998).

7 Rahmouni, K., Haynes, W. G., Morgan, D. A. & Mark, A. L. Intracellular Mechanisms Involved in Leptin Regulation of Sympathetic Outflow. Hypertension 41, 763–767, doi:10.1161/01.HYP.0000048342.54392.40 (2003).

8 Alawieh, A., Langley, E. F. & Tomlinson, S. Targeted complement inhibition salvages stressed neurons and inhibits neuroinflammation after stroke in mice. Science Translational Medicine 10, eaao6459, doi:10.1126/scitranslmed.aao6459 (2018).

9 Propson, N. E., Roy, E. R., Litvinchuk, A., Köhl, J. & Zheng, H. Endothelial C3a receptor mediates vascular inflammation and blood-brain barrier permeability during aging. J Clin Invest 131, doi:10.1172/jci140966 (2021).

10 Davoust, N., Jones, J., Stahel, P. F., Ames, R. S. & Barnum, S. R. Receptor for the C3a anaphylatoxin is expressed by neurons and glial cells. Glia 26, 201–211, doi:10.1002/(sici)1098-1136(199905)26:3<201::aid-glia2>3.0.co;2-m (1999).

11 Sun, R. et al. C3aR in the medial prefrontal cortex modulates the susceptibility to LPS-induced depressive-like behaviors through glutamatergic neuronal excitability. Prog Neurobiol 236, 102614, doi:10.1016/j.pneurobio.2024.102614 (2024).

12 Schadt, E. E. et al. An integrative genomics approach to infer causal associations between gene expression and disease. Nat Genet 37, 710–717, doi:10.1038/ng1589 (2005).

13 Mamane, Y. et al. The C3a anaphylatoxin receptor is a key mediator of insulin resistance and functions by modulating adipose tissue macrophage infiltration and activation. Diabetes 58, 2006–2017, doi:10.2337/db09-0323 (2009).

14 Feng, L., Zhao, Y. & Wang, W. L. Association between complement C3 and the prevalence of metabolic-associated fatty liver disease in a Chinese population: a cross-sectional study. BMJ Open 11, e051218, doi:10.1136/bmjopen-2021-051218 (2021).

15 Schupf, N., Williams, C. A., Hugli, T. E. & Cox, J. Psychopharmacological activity of anaphylatoxin C3a in rat hypothalamus. J Neuroimmunol 5, 305–316, doi:10.1016/0165-5728(83)90051-6 (1983).

16 Srakočić, S. et al. Proposed practical protocol for flow cytometry analysis of microglia from the healthy adult mouse brain: Systematic review and isolation methods’ evaluation. Frontiers in Cellular Neuroscience Volume 16 - 2022, doi:10.3389/fncel.2022.1017976 (2022).

17 Valdearcos, M. et al. Microglial Inflammatory Signaling Orchestrates the Hypothalamic Immune Response to Dietary Excess and Mediates Obesity Susceptibility. Cell Metab 26, 185–197.e183, doi:10.1016/j.cmet.2017.05.015 (2017).

18 Dapper, C. et al. The antiobese effect of AT1 receptor blockade is augmented in mice lacking Mas. Naunyn-Schmiedeberg’s archives of pharmacology 392, 865–877, doi:10.1007/s00210-019-01643-0 (2019).

19 Quell, K. M. et al. Monitoring C3aR Expression Using a Floxed tdTomato-C3aR Reporter Knock-in Mouse. J Immunol 199, 688–706, doi:10.4049/jimmunol.1700318 (2017).

20 Ghilardi, N. & Skoda, R. C. The leptin receptor activates janus kinase 2 and signals for proliferation in a factor-dependent cell line. Mol Endocrinol 11, 393–399, doi:10.1210/mend.11.4.9907 (1997).

21 Tang, J. et al. JR14a: A novel antagonist of C3aR attenuates neuroinflammation in cerebral ischemia-reperfusion injury. Brain Res Bull 213, 110986, doi:10.1016/j.brainresbull.2024.110986 (2024).

22 Rowley, J. A. et al. Potent Thiophene Antagonists of Human Complement C3a Receptor with Anti-Inflammatory Activity. J Med Chem 63, 529–541, doi:10.1021/acs.jmedchem.9b00927 (2020).

23 DeFalco, J. et al. Virus-assisted mapping of neural inputs to a feeding center in the hypothalamus. Science 291, 2608–2613, doi:10.1126/science.1056602 (2001).

24 Plum, L. et al. Enhanced leptin-stimulated Pi3k activation in the CNS promotes white adipose tissue transdifferentiation. Cell Metab 6, 431–445, doi:10.1016/j.cmet.2007.10.012 (2007).

25 Zhong, X. et al. Essential role of MFSD1-GLMP-GIMAP5 in lymphocyte survival and liver homeostasis. Proc Natl Acad Sci U S A 120, e2314429120, doi:10.1073/pnas.2314429120 (2023).

26 Packer, M., Schiattarella, G. G. & Borlaug, B. What Exactly Is Cardiometabolic HFpEF: A Phenotype or an Endotype? Circulation: Heart Failure 0, e014031, doi:10.1161/CIRCHEARTFAILURE.125.014031.

27 Daou, D., Tong, D., Schiattarella, G. G., Gillette, T. G. & Hill, J. A. What Is Cardiometabolic HFpEF and How Can We Study it Preclinically? JACC: Basic to Translational Science 10, 101295, 10.1016/j.jacbts.2025.04.009 (2025).

28 Zhao, S., Kusminski, C. M. & Scherer, P. E. Adiponectin, Leptin and Cardiovascular Disorders. Circulation Research 128, 136–149, doi:10.1161/CIRCRESAHA.120.314458 (2021).

29 Belin de Chantemèle, E. J., Mintz, J. D., Rainey, W. E. & Stepp, D. W. Impact of leptin-mediated sympatho-activation on cardiovascular function in obese mice. Hypertension 58, 271–279, doi:10.1161/hypertensionaha.110.168427 (2011).

30 Ursini, F. et al. Serum complement C3 strongly correlates with whole-body insulin sensitivity in rheumatoid arthritis. Clin Exp Rheumatol 35, 18–23 (2017).

31 Pomeroy, C. et al. Effect of body weight and caloric restriction on serum complement proteins, including Factor D/adipsin: studies in anorexia nervosa and obesity. Clin Exp Immunol 108, 507–515, doi:10.1046/j.1365-2249.1997.3921287.x (1997).

32 Horvath, T. L. et al. Synaptic input organization of the melanocortin system predicts diet-induced hypothalamic reactive gliosis and obesity. Proc Natl Acad Sci U S A 107, 14875–14880, doi:10.1073/pnas.1004282107 (2010).

33 Summers, S. A., Chaurasia, B. & Holland, W. L. Metabolic Messengers: ceramides. Nature Metabolism 1, 1051–1058, doi:10.1038/s42255-019-0134-8 (2019).

34 De Souza, C. T. et al. Consumption of a fat-rich diet activates a proinflammatory response and induces insulin resistance in the hypothalamus. Endocrinology 146, 4192–4199, doi:10.1210/en.2004-1520 (2005).

35 Zhou, Y. & Rui, L. Leptin signaling and leptin resistance. Front Med 7, 207–222, doi:10.1007/s11684-013-0263-5 (2013).

36 Contreras, C. et al. Central ceramide-induced hypothalamic lipotoxicity and ER stress regulate energy balance. Cell Rep 9, 366–377, doi:10.1016/j.celrep.2014.08.057 (2014).

37 Zabolotny, J. M. et al. Protein-tyrosine phosphatase 1B expression is induced by inflammation in vivo. J Biol Chem 283, 14230–14241, doi:10.1074/jbc.M800061200 (2008).

38 Zhang, X. et al. Hypothalamic IKKbeta/NF-kappaB and ER stress link overnutrition to energy imbalance and obesity. Cell 135, 61–73, doi:10.1016/j.cell.2008.07.043 (2008).

39 Schiattarella, G. G. et al. Nitrosative stress drives heart failure with preserved ejection fraction. Nature 568, 351–356, doi:10.1038/s41586-019-1100-z (2019).

40 Tadinada, S. M. et al. Functional resilience of C57BL/6J mouse heart to dietary fat overload. American Journal of Physiology-Heart and Circulatory Physiology 321, H850–H864, doi:10.1152/ajpheart.00419.2021 (2021).

41 Manrique, C. et al. Obesity and insulin resistance induce early development of diastolic dysfunction in young female mice fed a Western diet. Endocrinology 154, 3632–3642, doi:10.1210/en.2013-1256 (2013).

42 Carbone, S. et al. A high-sugar and high-fat diet impairs cardiac systolic and diastolic function in mice. Int J Cardiol 198, 66–69, doi:10.1016/j.ijcard.2015.06.136 (2015).

43 Bostick, B. et al. Mineralocorticoid receptor blockade prevents Western diet-induced diastolic dysfunction in female mice. American Journal of Physiology-Heart and Circulatory Physiology 308, H1126–H1135, doi:10.1152/ajpheart.00898.2014 (2015).

44 Maurya Santosh, K., Carley Andrew, N., Maurya Chandan, K. & Lewandowski, E. D. Western Diet Causes Heart Failure With Reduced Ejection Fraction and Metabolic Shifts After Diastolic Dysfunction and Novel Cardiac Lipid Derangements. JACC: Basic to Translational Science 8, 422–435, doi:10.1016/j.jacbts.2022.10.009 (2023).

45 Moreno-Navarrete, J. M. & Fernández-Real, J. M. The complement system is dysfunctional in metabolic disease: Evidences in plasma and adipose tissue from obese and insulin resistant subjects. Seminars in Cell & Developmental Biology 85, 164–172, 10.1016/j.semcdb.2017.10.025 (2019).

46 Hernández-Mijares, A. et al. Effect of weight loss on C3 and C4 components of complement in obese patients. European Journal of Clinical Investigation 42, 503–509, 10.1111/j.1365-2362.2011.02606.x (2012).

47 Kosiborod Mikhail, N., et al. Semaglutide in Patients with Heart Failure with Preserved Ejection Fraction and Obesity. New England Journal of Medicine 389, 1069–1084, doi:10.1056/NEJMoa2306963 (2023).

48 Kosiborod Mikhail, N., et al. Semaglutide in Patients with Obesity-Related Heart Failure and Type 2 Diabetes. New England Journal of Medicine 390, 1394–1407, doi:10.1056/NEJMoa2313917 (2024).

49 Lee, A. A. et al. Incretin-Based Therapies Through the Decades: Molecular Innovations and Clinical Impact. Medical Sciences 13, 269 (2025).

50 Milanski, M. et al. Inhibition of hypothalamic inflammation reverses diet-induced insulin resistance in the liver. Diabetes 61, 1455–1462, doi:10.2337/db11-0390 (2012).

51 Desai, J. V. et al. C5a-licensed phagocytes drive sterilizing immunity during systemic fungal infection. Cell 186, 2802–2822.e2822, doi:10.1016/j.cell.2023.04.031 (2023).

52 He, S. et al. A complement-dependent balance between hepatic ischemia/reperfusion injury and liver regeneration in mice. J Clin Invest 119, 2304–2316, doi:10.1172/jci38289 (2009).

53 Wysoczynski, M. et al. Defective engraftment of C3aR-/-hematopoietic stem progenitor cells shows a novel role of the C3a-C3aR axis in bone marrow homing. Leukemia 23, 1455–1461, doi:10.1038/leu.2009.73 (2009).

54 Bartolomucci, A. et al. TLQP-21, a VGF-derived peptide, increases energy expenditure and prevents the early phase of diet-induced obesity. Proc Natl Acad Sci U S A 103, 14584–14589, doi:10.1073/pnas.0606102103 (2006).

55 Luo, P. et al. Structural insights into the agonist activity of the nonpeptide modulator JR14a on C3aR. Cell Discovery 11, 7, doi:10.1038/s41421-024-00765-x (2025).

56 Kim, J. et al. Structural insights into small-molecule agonist recognition and activation of complement receptor C3aR. Embo j 44, 2803–2826, doi:10.1038/s44318-025-00429-w (2025).

57 Li, F. & Cui, S. Knockdown of C3aR alleviates age-related bone loss *via* activation of YAP1/β-catenin signaling. Journal of Biological Chemistry 301, doi:10.1016/j.jbc.2025.108500 (2025).

58 Virtue, S. & Vidal-Puig, A. GTTs and ITTs in mice: simple tests, complex answers. Nat Metab 3, 883–886, doi:10.1038/s42255-021-00414-7 (2021).

59 Even, P. C. & Nadkarni, N. A. Indirect calorimetry in laboratory mice and rats: principles, practical considerations, interpretation and perspectives. Am J Physiol Regul Integr Comp Physiol 303, R459–476, doi:10.1152/ajpregu.00137.2012 (2012).

