## Extended Data for "C3a receptor signalling in ObRb-expressing neurons promotes leptin resistance and obesity-associated cardiac remodelling"

#### Extended Data Figures and Supplementary Tables

#### Extended Data Figure 1

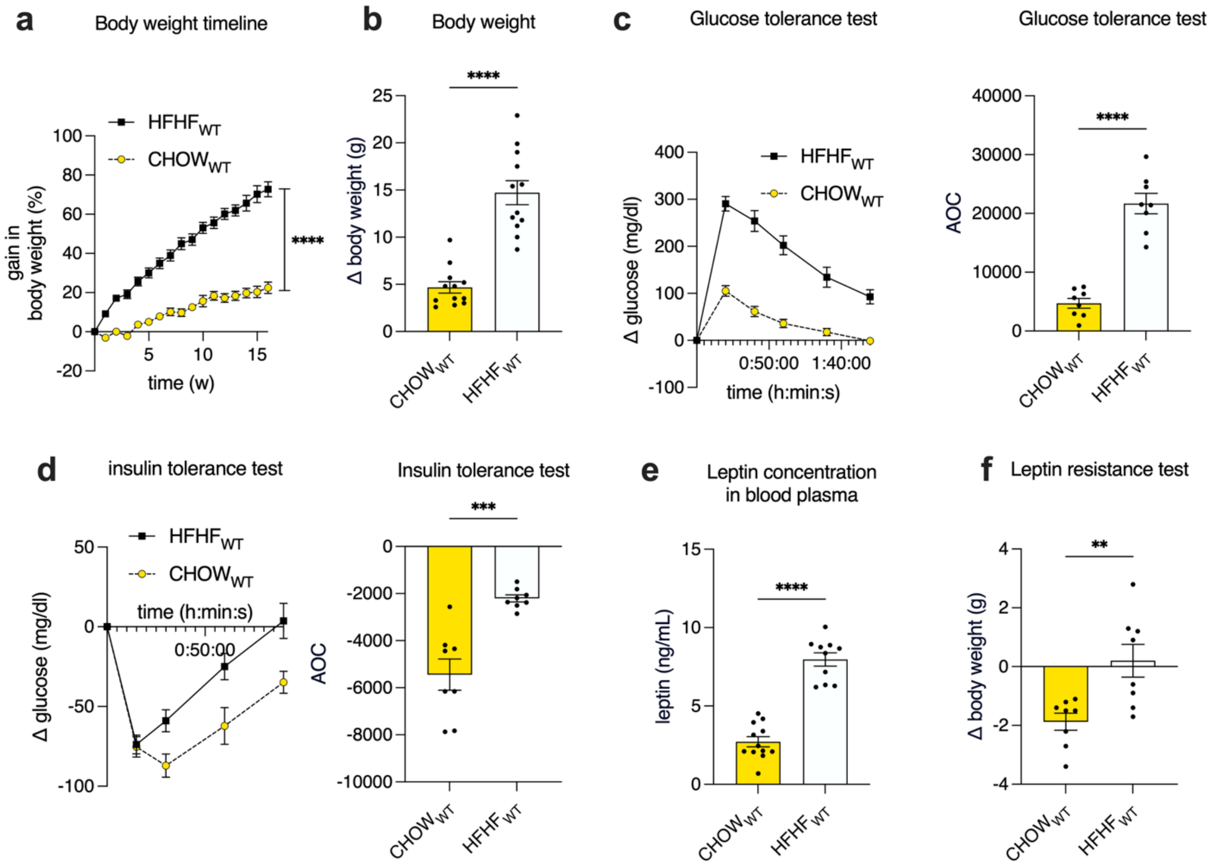

**Extended Data Fig. 1. Impact of high-fat/high-fructose diet on body weight and metabolic parameters. a,** Relative body weight gain of WT mice fed a high-fat/high-fructose (HFHF) diet or control diet (CHOW) over 16 weeks, expressed as percentage change from baseline weight (n=12/group). **b,** Body weight gain in WT mice 16 weeks after HFHF or CHOW relative to baseline weight (n=12/group). **c, Left:** Baseline-subtracted glucose levels during glucose tolerance test (GTT) at week 12 (n = 8). **Right**: Area over the curve (AOC) from GTT calculated from baseline-subtracted glucose levels (n=8). **d, Left:** Baseline-subtracted glucose levels during insulin tolerance test (ITT) at week 13 (n= 8/group). **Right:** AOC from ITT calculated from baseline-subtracted glucose levels (n= 8/group). **e,** Leptin plasma levels measured by ELISA at week 16 (n=12 CHOW, n=10 HFHF). **f,** Body weight change assessed the day after the final leptin injection in the leptin resistance test (LRT) at week 14 (n=8/group). Data are shown as mean ± SEM. Statistical differences between groups were determined using two-way ANOVA (**a**) or unpaired Student's t-test (**b-f**). *$p<0.05$, **$p<0.01$, ***$p<0.001$, ****$P<0.0001$.

#### Extended Data Figure 2

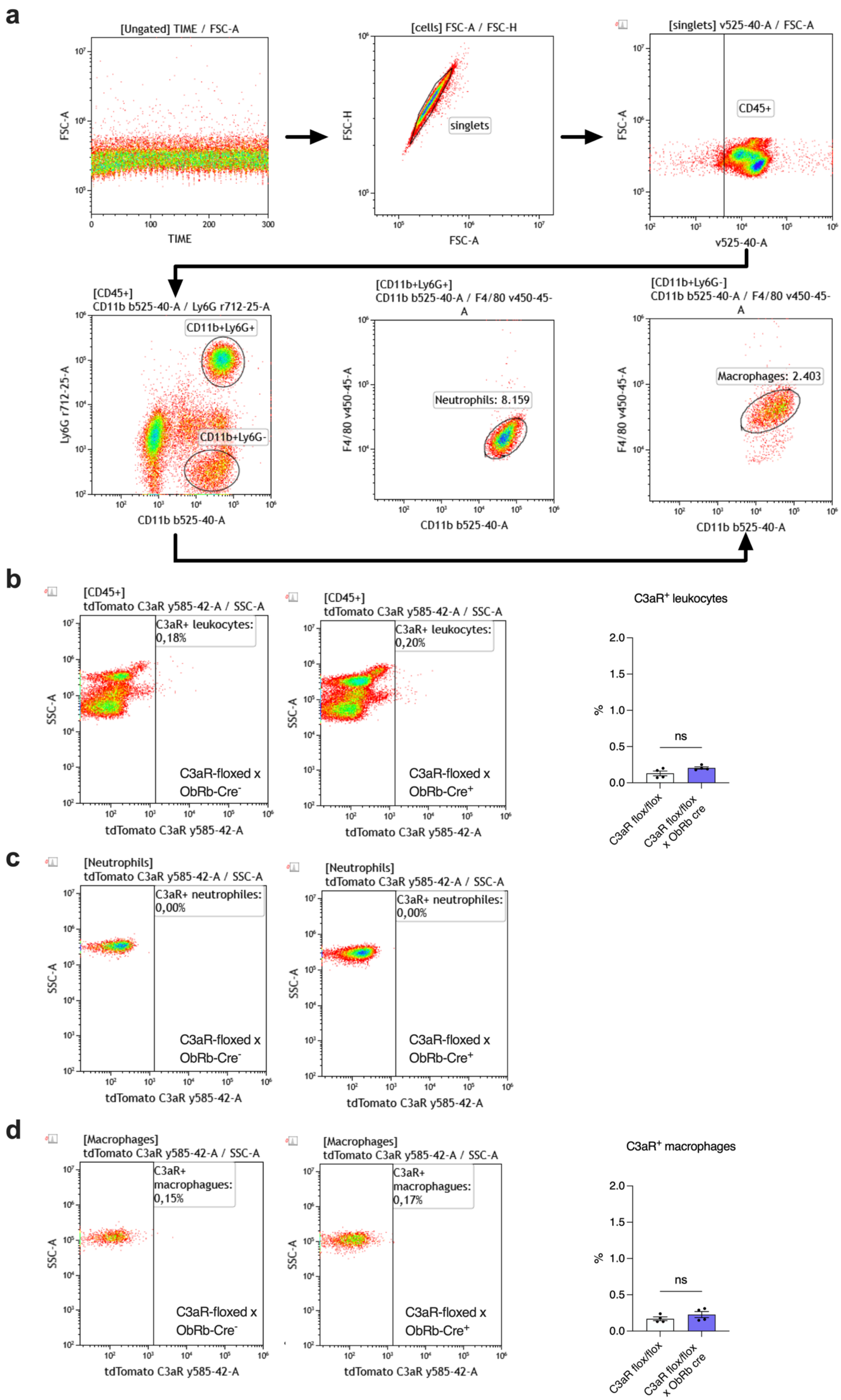

**Extended Data Fig. 2. Flow cytometric analysis of C3aR expression on murine blood leukocytes from C3aR$^{flox/flox}$ ObRb-Cre$^{+}$ mice and ObRb-Cre$^{-}$ littermates**. **a**, Gating strategy to identify CD11b$^{+}$Ly6G$^{+}$ neutrophils and CD11b$^{+}$Ly6G$^{-}$F4/80$^{+}$ macrophages. **b-d, Left and Middle:** Dot-plot showing tdTomato expression in CD45$^{+}$ leukocytes (**b**), CD11b$^{+}$Ly6G$^{+}$ neutrophils (**c**), and CD11b$^{+}$Ly6G$^{-}$F4/80$^{+}$ macrophages (**d**) from naive C3aR$^{flox/flox}$ ObRb-Cre$^{+}$ mice and ObRb-Cre$^{-}$ littermates. **Right:** Frequency of CD45$^{+}$ leukocytes (**b**) and CD11b$^{+}$Ly6G$^{-}$F4/80$^{+}$ macrophages (**d**) from C3aR$^{flox/flox}$ ObRb-Cre$^{+}$ and ObRb-Cre$^{-}$ littermates expressing tdTomato. (n=4 per group). Data shown are the mean ± SEM. Statistical differences between groups were assessed using Mann–Whitney U-test; ns: not significant.

### Extended Data Figure 3

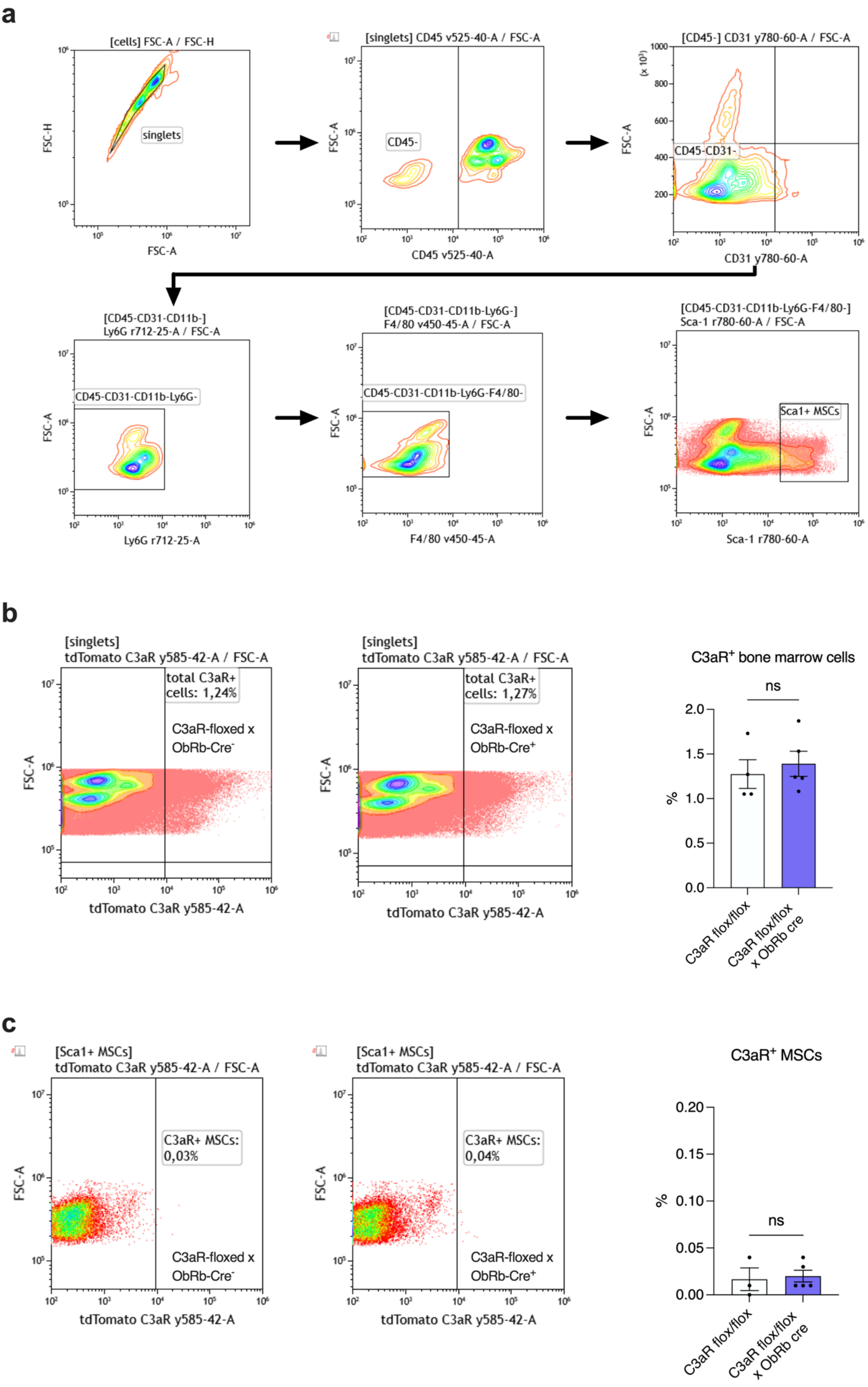

**Extended Data Fig. 3. Flow cytometric analysis of C3aR expression on bone marrow-derived CD45$^{+}$ leukocytes and CD45$^{-}$Sca$^{+}$ mesenchymal stem cells (MSCs) from C3aR$^{flox/flox}$ ObRb-Cre$^{+}$ mice and ObRb-Cre$^{–}$ littermates**. **a**, Gating strategy to identify CD45$^{-}$CD31$^{-}$CD11b$^{-}$Ly6G$^{-}$Sca$^{+}$MSCs. **b-c, Left and Middle:** Dot-plot showing tdTomato expression in CD45$^{+}$ leukocytes (**b**) and CD45$^{-}$CD31$^{-}$CD11b$^{-}$Ly6G$^{-}$Sca$^{+}$MSCs (**c**). **Right:** Frequency of CD45$^{+}$ leukocytes (**b**) and CD45$^{-}$CD31$^{-}$CD11b$^{-}$Ly6G$^{-}$Sca$^{+}$MSCs (**c**) leukocytes from C3aR$^{flox/flox}$ ObRb-Cre$^{+}$ and ObRb-Cre$^{–}$ littermates expressing tdTomato. (n=4 per group). Data shown are the mean ± SEM. Statistical differences between groups were assessed using Mann–Whitney U-test; ns: not significant.

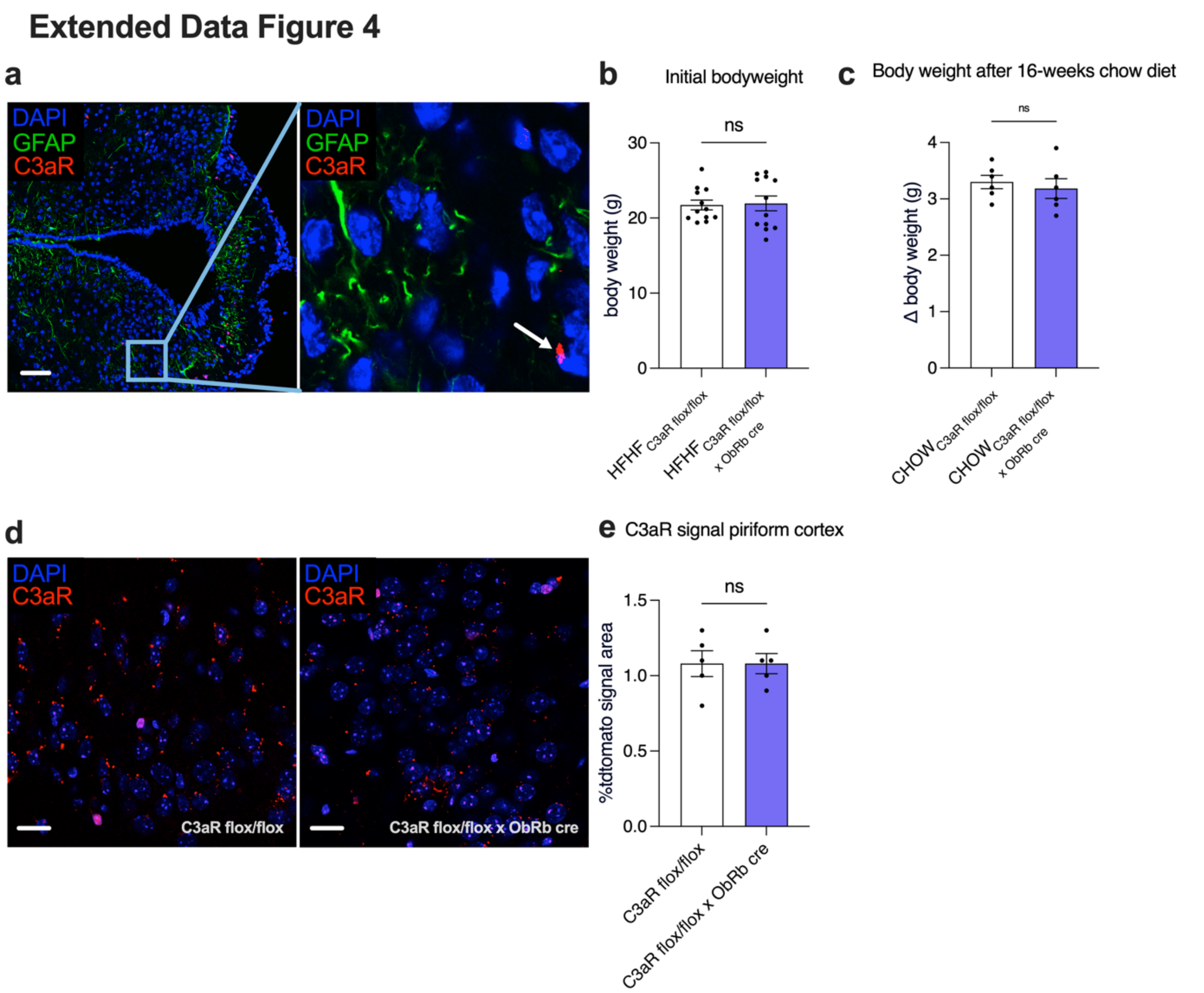

**Extended Data Fig. 4. ObRb-Cre does not detectably alter tdTomato-C3aR reporter signal in hypothalamic astrocytes or piriform cortex and does not affect body weight under chow. a,** Representative immunofluorescence image of hypothalamic sections from tdTomato-labeled C3aR$^{flox/flox}$ mice crossed with ObRb-Cre mice, showing co-staining for GFAP (green, astrocytes), tdTomato-C3aR (red), and DAPI (blue). No co-localization of C3aR with GFAP-positive astrocytes in hypothalamic sections was observed. The white arrows point toward C3aR$^{+}$ cells that do not express GFAP. Scale bar: 100 µm. **b,** Initial body weight of C3aR$^{flox/flox}$ and C3aR$^{flox/flox}$ x ObRb-Cre mice. No significant differences were observed between groups prior to the start of HFHF (n=12/group). **c,** Body weight gain of C3aR$^{flox/flox}$ and C3aR$^{flox/flox}$ x ObRb-Cre mice after 12 weeks of chow diet (n=6/group). **d,** Representative immunofluorescence image of the piriform cortex showing tdTomato-labelled C3aR expression (red) with DAPI nuclear staining (blue). Scale bar, 20 µm. **e,** Quantification of tdTomato-positive C3aR signals in C3aR$^{flox/flox}$ and C3aR$^{flox/flox}$ x ObRb-Cre mice in the piriform cortex. (n=5/group). All data are shown as mean ± SEM. Statistical analyses were performed using an unpaired Student's t-test (**b, c, e**); ns: not significant.

#### Extended Data Figure 5

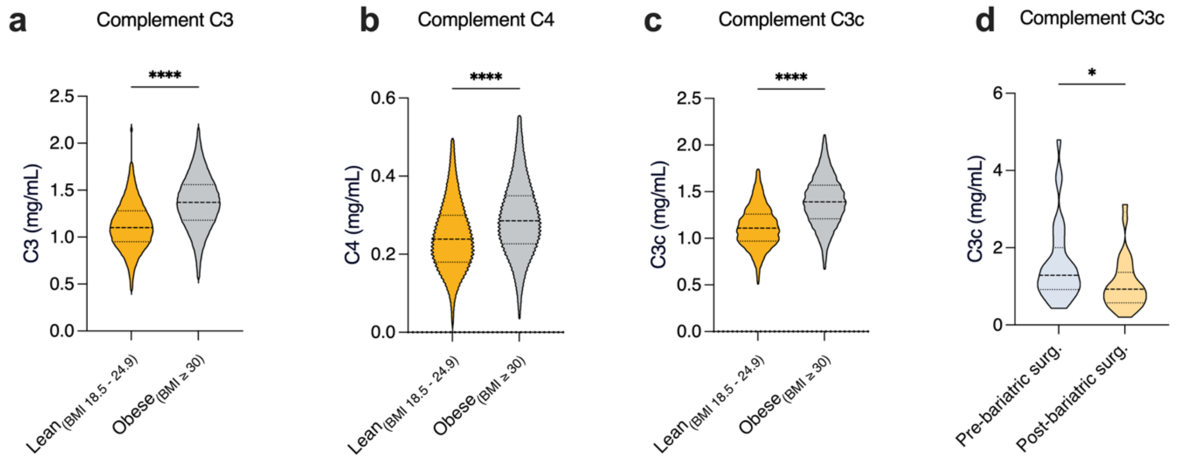

**Extended Data Fig. 5. Complement activation in human obesity.** Violin plots of complement activation in obese humans. **a–c**, Plasma levels of complement factors C3 (**a**), C4 (**b**) and the C3 cleavage fragment C3c (**c**) in humans with normal BMI (18.5 – 24.9 kg/m², n=268,769) and obesity (BMI >30 kg/m², n=291,148). **d**, C3c plasma levels in obese patients before and 12 months after bariatric surgery (n=56). Statistical analyses were performed using an unpaired Student's t-test (a-c) and paired t-test (d). *p<0.05, ****p<0.0001.

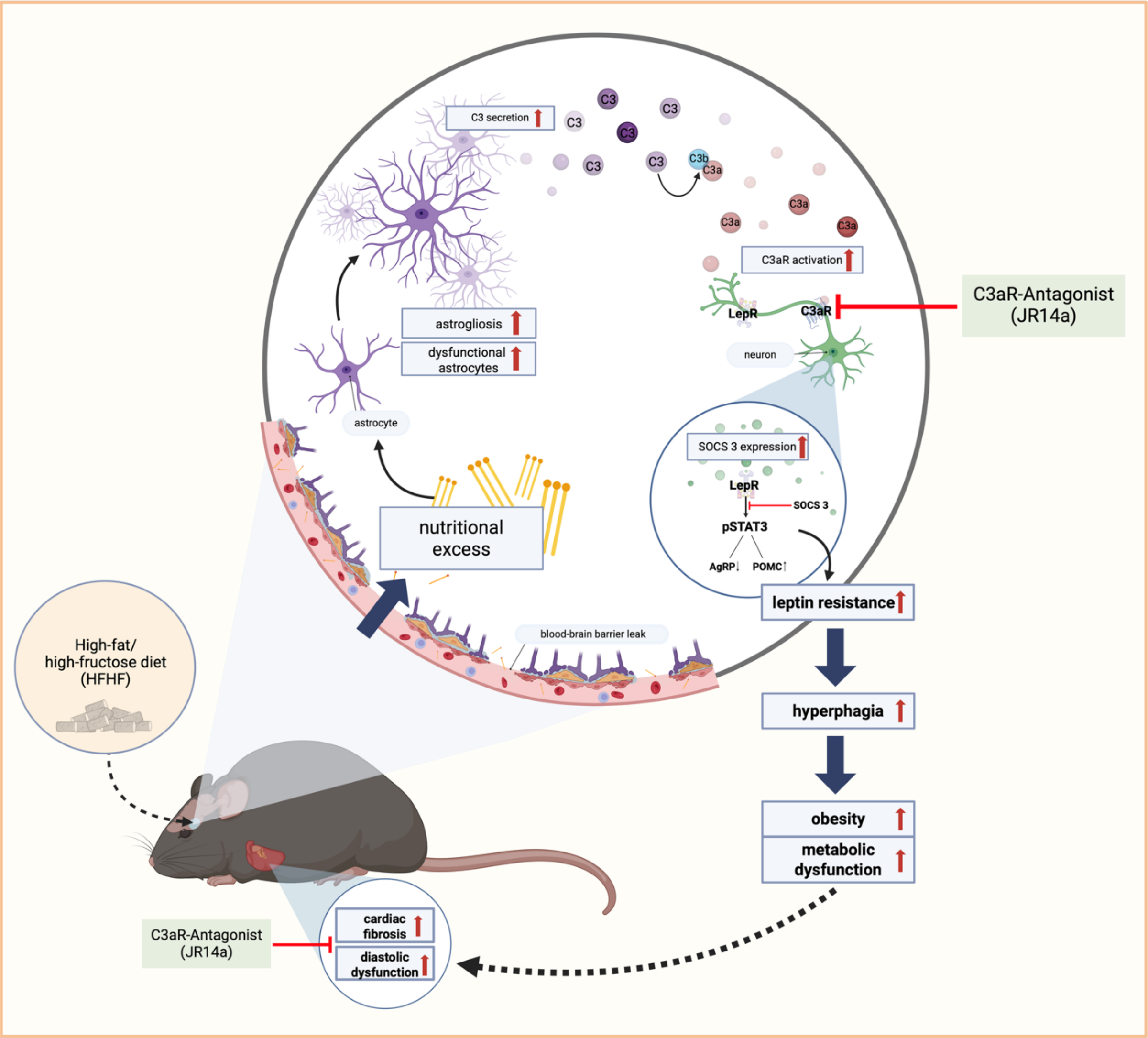

**Extended Data Fig. 6.** Dietary excess is associated with hypothalamic gliosis and local complement activation, including increased C3a and activated C3 fragments. Locally generated C3a may act on C3a receptors on leptin-responsive hypothalamic neurons to promote SOCS3 expression and impair leptin signalling. Genetic deletion or pharmacological targeting of C3aR improves leptin responsiveness during HFHF feeding and attenuates obesity-associated metabolic and cardiac remodelling. Together, these data support a glia-to-neuron complement signalling pathway contributing to hypothalamic leptin resistance and obesity-associated diastolic impairment.

| Tissue | Group | *C3ar1* | *Cdkn1b* | delta_Ct | rel_expression |
|---|---|---|---|---|---|
| **Gonadal White Adipose Tissue** | Cre+ | 36.092 | 22.470 | 13.62196159 | 7.93E-05 |
| **Gonadal White Adipose Tissue** | Cre+ | not detectable | 23.838 | not detectable | not detectable |
| **Gonadal White Adipose Tissue** | Cre- | 38.399 | 22.395 | 16.00389845 | 1.52E-05 |
| **Gonadal White Adipose Tissue** | Cre+ | 35.890 | 22.175 | 13.71529545 | 7.44E-05 |
| **Gonadal White Adipose Tissue** | Cre+ | 37.523 | 22.316 | 15.20695519 | 2.64E-05 |
| **Gonadal White Adipose Tissue** | Cre+ | 37.029 | 22.472 | 14.55688544 | 4.15E-05 |
| **Gonadal White Adipose Tissue** | Cre- | 38.728 | 22.396 | 16.33205445 | 1.21E-05 |
| **Gonadal White Adipose Tissue** | Cre- | 36.694 | 22.364 | 14.33069134 | 4.85E-05 |
| **Gonadal White Adipose Tissue** | Cre- | 36.177 | 22.804 | 13.37273026 | 9.43E-05 |
| **Brown Adipose Tissue** | Cre+ | 38.913 | 23.217 | not detectable | not detectable |
| **Brown Adipose Tissue** | Cre+ | not detectable | 22.808 | not detectable | not detectable |
| **Brown Adipose Tissue** | Cre- | not detectable | 23.099 | not detectable | not detectable |
| **Brown Adipose Tissue** | Cre+ | 38.054 | 22.484 | not detectable | not detectable |
| **Brown Adipose Tissue** | Cre+ | not detectable | 22.705 | not detectable | not detectable |
| **Brown Adipose Tissue** | Cre+ | 36.438 | 22.278 | not detectable | not detectable |
| **Brown Adipose Tissue** | Cre- | not detectable | 22.649 | not detectable | not detectable |
| **Brown Adipose Tissue** | Cre- | 36.582 | 23.088 | not detectable | not detectable |
| **Brown Adipose Tissue** | Cre- | not detectable | 23.137 | not detectable | not detectable |
| **Liver** | Cre+ | not detectable | 23.978 | not detectable | not detectable |
| **Liver** | Cre+ | not detectable | 23.985 | not detectable | not detectable |
| **Liver** | Cre- | not detectable | 24.302 | not detectable | not detectable |
| **Liver** | Cre+ | not detectable | 23.420 | not detectable | not detectable |
| **Liver** | Cre+ | not detectable | 23.630 | not detectable | not detectable |
| **Liver** | Cre+ | not detectable | 23.225 | not detectable | not detectable |
| **Liver** | Cre- | not detectable | 23.900 | not detectable | not detectable |
| **Liver** | Cre- | not detectable | 24.031 | not detectable | not detectable |
| **Liver** | Cre- | not detectable | 23.820 | not detectable | not detectable |
| **Pancreas** | Cre+ | not detectable | 28.789 | not detectable | not detectable |
| **Pancreas** | Cre+ | not detectable | 28.613 | not detectable | not detectable |
| **Pancreas** | Cre- | not detectable | 27.774 | not detectable | not detectable |
| **Pancreas** | Cre+ | not detectable | 26.107 | not detectable | not detectable |
| **Pancreas** | Cre+ | not detectable | 27.435 | not detectable | not detectable |
| **Pancreas** | Cre+ | not detectable | 27.256 | not detectable | not detectable |
| **Pancreas** | Cre- | not detectable | 27.514 | not detectable | not detectable |
| **Pancreas** | Cre- | not detectable | 29.777 | not detectable | not detectable |
| **Pancreas** | Cre- | not detectable | 28.855 | not detectable | not detectable |
| **Heart** | Cre+ | 35.910 | 24.411 | 11.499 | 3.46E-04 |
| **Heart** | Cre+ | 36.209 | 23.945 | 12.264 | 2.03E-04 |
| **Heart** | Cre- | 37.584 | 24.290 | 13.294 | 9.96E-05 |

| | | | | | |
|---|---|---|---|---|---|
| **Heart** | Cre+ | 36.850 | 23.301 | 13.549 | 8.34E-05 |
| **Heart** | Cre+ | 37.392 | 24.405 | 12.987 | 1.23E-04 |
| **Heart** | Cre+ | not detectable | 23.451 | not detectable | not detectable |
| **Heart** | Cre- | 35.728 | 23.648 | 12.08 | 2.31E-04 |
| **Heart** | Cre- | 36.462 | 24.549 | 11.913 | 2.59E-04 |
| **Heart** | Cre- | 37.455 | 24.415 | 13.04 | 1.19E-04 |

**Supplementary Table 1: Quantitative evaluation of *C3ar1* gene expression in different visceral tissues of tdTomato-C3aR$^{flox/flox}$ ObRb-Cre$^{+}$ and tdTomato-C3aR$^{flox/flox}$ x ObRb-Cre$^{-}$ mice**. Shown are Ct values and the relative gene expression of *C3ar1* normalized to *Cdkn1b* (ΔCt and 2^–ΔCt) in the indicated tissues of Cre-expressing and non-Cre–expressing tdTomato-C3aR$^{flox/flox}$ mice (n=4-5/group).

| | Lean (n=268,769) | Obese (n=291,148) | P-Value |
|---|---|---|---|
| Age | 56.2 ± 20.5 | 56.7 ± 16.7 | < 0.0001 |
| Female (%) | 68 % | 68 % | |
| BMI (kg/m²) | 22.1 ± 1.76 | 36.5 ± 6.2 | < 0.0001 |
| Type 2 diabetes mellitus (%) | 17 | 34 | < 0.0001 |
| Hypertensive disease (%) | 43 | 62 | < 0.0001 |
| Hyperlipidemia (%) | 36 | 49 | < 0.0001 |
| Coronary Heart Disease (%) | 21 | 24 | < 0.0001 |
| Heart failure (%) | 16 | 20 | < 0.0001 |
| Chronic kidney disease (%) | 26 | 30 | < 0.0001 |

**Supplementary Table 2. Baseline clinical characteristics in obese and lean individuals from the TriNetX cohort**

| | Patients undergoing bariatric operation |
|---|---|
| Age | 43.3 ± 10.6 |
| Female (%) | 83.3 |
| BMI (kg/m$^2$) pre-surgery | 47.4 ± 6.5 |
| BMI (kg/m$^2$) 12 months post-surgery | 31.7 ± 5.5 |
| Type 2 diabetes mellitus (%) | 22 |
| Hypertensive disease (%) | 53 |
| Hyperlipidemia (%) | 14.7 |
| Obstructive sleep apnea (%) | 49 |
| Smoking (%) | 21 |

**Supplementary Table 3. Clinical characteristics of patients undergoing bariatric operation**
